# Identifying potential biomarkers of response to CCR5Δ32 HSCT in HIV infection

**DOI:** 10.1101/2025.03.30.646188

**Authors:** Jason Chang, Feilim Mac Gabhann

## Abstract

While HIV can be effectively suppressed to a chronic, mostly asymptomatic infection with combination antiretroviral therapy (cART), a cure is still needed. Hematopoietic stem cell transplantation (HSCT) from HIV-resistant donors has shown promise and has resulted in HIV remission in five patients. However, this treatment strategy does not guarantee HIV remission; six other patients who received a similar transplant had poor outcomes and died within a year of treatment. These different outcomes may be due to inter-individual differences in HIV infection dynamics that result in heterogeneity of therapeutic responses to HSCT. Using a previously published mechanistic model of HIV infection and virtual populations calibrated from patient data, we performed simulations to understand how different parameters in the model can influence the observed heterogeneity in therapeutic outcomes across virtual patient populations. Our simulations confirmed that discontinuation of cART, without HSCT, always leads to viral rebound, and that time to rebound differs across patients due to the interindividual variability (IIV) in underlying infection dynamics. Extending the duration of cART only slightly increased the predicted median time to rebound and its variance. By contrast, HSCT followed by cART cessation led to HIV remission, but only for a subset of the virtual patients. The proportion of patients predicted to go into remission depends directly on the ratio of donor to host cell immune cells in the post-HSCT chimeric immune system. Of the mechanistic model parameters, no single parameter determined whether a patient was a responder or a non-responder; rather, the interactions between multiple model parameters were crucial in driving treatment responses. In contrast, virtual equivalents of clinically accessible observations, e.g. viral load and cell populations at specific times, were shown to be better predictors than mechanistic model parameters in separating patients into non-responding (viral rebound) and responding (no rebound) clusters.

**One Sentence Summary:** Simulations using a mechanistic HIV model suggest that interindividual variability (IIV) in infection dynamics drives differential responses to hematopoietic stem cell transplantation (HSCT), with HIV remission being largely dependent on donor-to-host immune cell ratios in the post-HSCT chimeric immune system and better predicted by clinically accessible observations than individual model parameters.

## Introduction

HIV can be effectively suppressed and maintained as a chronic infection without progression to major disease by treatment with combination antiretroviral therapy (cART), a cocktail of drugs administered to suppress viral replication ^1^. However, cART is only a treatment and not a cure. Even after being on cART for years, patients will inevitably experience HIV rebound if they cease treatment ^2^, though there is heterogeneity in the time to viral rebound ^3^. The heterogeneity stems from the differences in the populations of various immune cells and the rates of various cellular processes that govern the infection dynamics within the patient.

To date, five patients have achieved HIV remission after the cessation of cART, and in all cases this followed a stem cell transplant, which suggests that this approach could be the basis for a HIV cure ^4^. Specifically, the common treatment was a transplantation of hematopoietic stem cells from donors that are homozygous for a mutation (Δ32 deletion) in the CC chemokine receptor 5 (CCR5) gene ^4^. The process by which HIV gains entry into host target cells involves the initial binding of the viral envelope glycoprotein (gp120) to its primary receptor, CD4, on the target cell surface. This is followed by the interactions with co-receptors CCR5 or C-X-C chemokine receptor 4 (CXCR4) ^5^. Given that the CCR5 gene plays a critical role in facilitating viral entry by some HIV subtypes, the CCR5Δ32 immune cells generated from these CCR5Δ32 stem cells are resistant to viral entry, do not get infected, and thus do not generate additional infectious progeny virions, resulting in a reduction in the overall viral burden in the transplant recipient ^6^. However, HSCT combined with cART does not guarantee HIV remission; six patients who have received allogeneic transplantation of CCR5-deficient stem cells had poor outcomes and died within a year of HSCT^7^.

Our laboratory previously published a mechanistic model of HIV dynamics to simulate the outcome of HSCT transplant, including both wild type and CCR5Δ32 donors ^8^. Using a system of ordinary differential equations, the model simulates the changes in viral load and the levels of various immune cell subpopulations over time. The simulation encompasses the entire infection timeline, from the initial onset of HIV infection, through a succession of treatment periods (or lack thereof), allowing for simulations of complex virtual clinical trials *in silico*. The model was parameterized using patient data from the Multicenter AIDS Cohort Study, under which the approximate dates of HIV seroconversion and AIDS diagnosis were available ^8^. To mimic the interindividual variability (IIV) in the HIV patient population, four virtual populations were created using the measured patient data for viral loads and T cell populations over time. Individual virtual patients are defined both by their initial conditions (levels of various immune cell subpopulations before initial exposure to HIV) and by the model parameters governing the dynamics of their immune system and the infection. In that previous study, the medians and 5th-95th percentile ranges of the virtual populations were validated to match those of the clinical data. The virtual patients across the four populations were categorized based on the time from HIV seroconversion to AIDS diagnosis in the absence of treatment.

Using this established mechanistic model of HIV dynamics and virtual populations, the motivation for this current study is to understand how the different parameters in the model determine whether a virtual patient responds or does not respond positively to HSCT; and also how the parameters may drive the observed heterogeneity in time to viral rebound following the cessation of cART.

## Materials and Methods

### Model description

The simulations performed in this study were based on a previously published mechanistic model of HIV dynamics ^8^. The model is a system of ordinary differential equations (ODEs) based on the processes affecting the dynamics of viral load and the levels of various immune cell subpopulations. Using these equations, we can simulate the time course of infection as well as multiple treatments including cART and HSCT, across multiple years. Key elements in the model include: uninfected CD4+ T cells and macrophages; latently infected and productively infected CD4+ T cells; infected macrophages; cytotoxic CD8+ T cells; and the virus itself (Figure 1). Treatments simulated include cART and HSCT. In the case of HSCT, both wild-type and augmented immune cell subpopulations are included in the model to reflect the chimeric nature of the immune system post-transplant. Each of these cell or virus populations is represented by an ODE, comprising rate terms that represent the biological processes that alter the state or abundance (e.g. cell production, cell death, virus infecting cell); each of these rate terms has model parameters that may be different from person to person. The model simulations were implemented and solved using MATLAB (MathWorks, Natick, MA)

**Figure 1.**
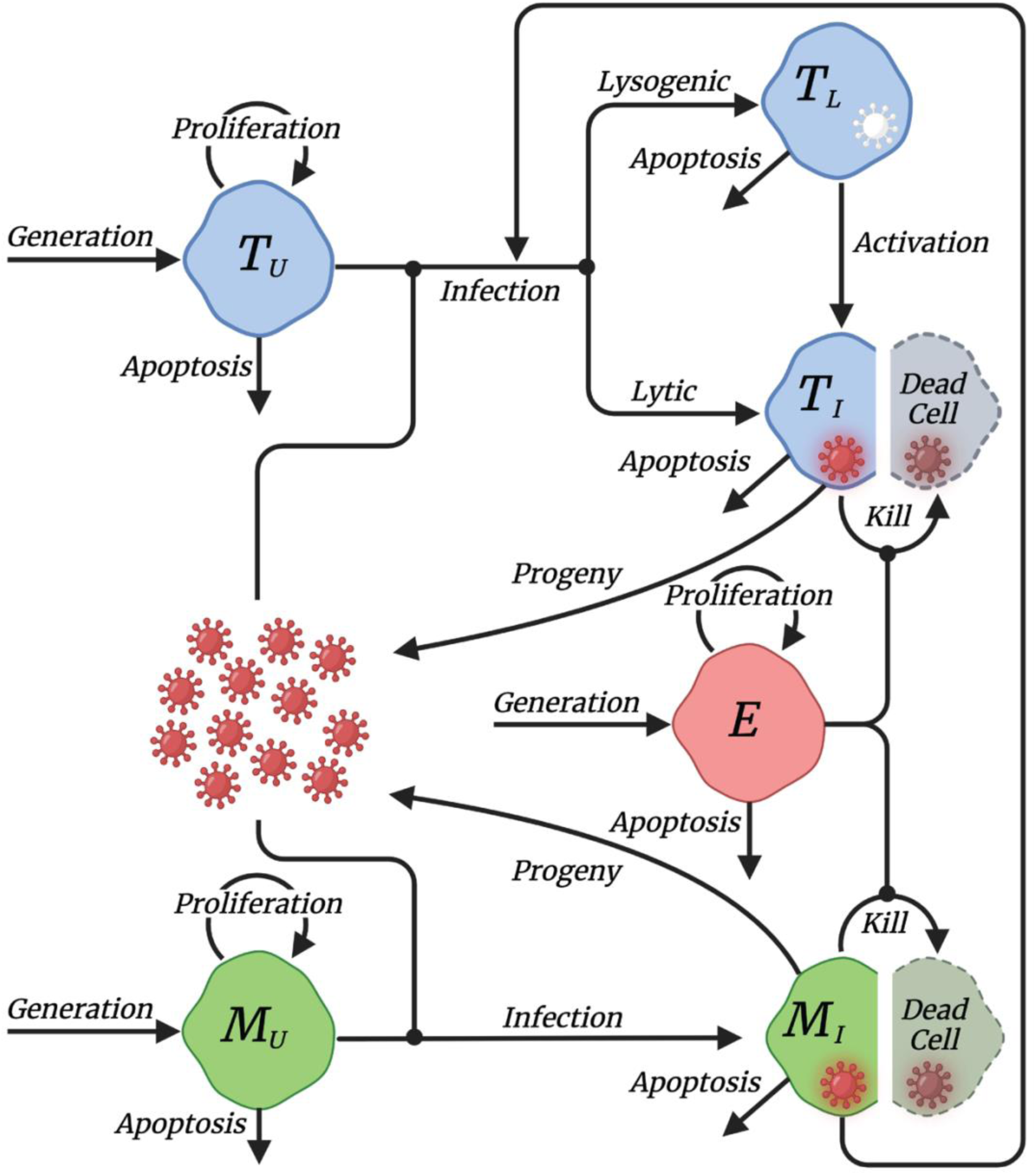
Mechanistic model of HIV infection. This is based on our previously-published model of HIV infection with wild-type and augmented immune cells under hematopoietic stem cell transplantation (HSCT) (Hosseini et al., 2016). The model is ODE-based and consists of key species that are involved in the infection dynamics: CD4+ T cells (T), macrophages (M), CD8+ T cells (E), and the virus (red particles). Under HSCT, the model also includes alternate versions of the cells shown to track both the wild-type immune cells and the augmented CD4+ T cells and macrophages, which carry the anti-HIV CCR5Δ32 mutation. The directed edges in the graph represent the processes that govern the level of different cell subpopulations and viruses.

### Generation of parameters for virtual populations

To capture the interindividual variability (IIV) in infection dynamics between HIV patients, we use the same four virtual patient populations created for the previous study ^8^. The virtual patients were categorized into four separate populations based on the time from HIV seroconversion to AIDS diagnosis when untreated: less than 3.5 years (rapid progressors, Population 1), between 3.5 and 7 years (Population 2), between 7 and 9 years (Population 3), and more than 9 years (slow progressors, Population 4). The virtual patient populations were generated based on patient viral loads and T cell population dynamics from real patient data for each category, with the medians and 5th-95th percentile ranges of the virtual population matching those of the clinical data ^8^. A total of 1000 virtual patients were generated for each population. Each virtual patient is defined by their initial conditions (levels of various immune cell subpopulations) as well as model parameters that define the dynamic response to infection.

### Simulation of HIV infection, treatment, and treatment interruption

We assumed the same level of initial viral exposure (V_0_ = 10^-3^ virions/μL) as the starting infection in our virtual patients. The initial levels of the immune cell subpopulations are patient-specific. The cART-mediated suppression of viral replication is reflected in the simulation by setting the infection rates of virus on uninfected CD4+ T cells (*k_VT_*), infected macrophages on uninfected CD4+ T cells (*k_MT_*), and virus on uninfected macrophages (*k_VM_*) all to zero. For treatment interruption (TI), i.e. when the treatment with cART is stopped, the infection rates are all assumed to return to their baseline values. The hematopoietic stem cell transplantation (HSCT) procedure takes advantage of *ex vivo* expansion of donor hematopoietic stem cells (HSCs) carrying the homozygous CCR5Δ32 mutation that confers HIV resistance. The process simulated in our model consists of total body irradiation (TBI) to suppress host immunity, followed by intravenous infusion and engraftment of the augmented donor HSCs. During TBI, irradiation of the bone marrow results in killing of both uninfected and productively infected CD4+ T cells, depletion of CD8+ T cells, and reduction of latent viral reservoir by two-thirds ^8–10^. Meanwhile, the tissue-residing macrophage subpopulations are mostly unaffected by the TBI treatment ^11^. However, TBI is not effective in completely eliminating the wild-type HSCs. Consequently, the host then rebuilds a chimeric immune system consisting of both wild-type and augmented immune cells following the infusion and engraftment of the donor cells. In our model, the level of chimerism is reflected in the production rates of uninfected CD4+ T cells and macrophages (*s_T_* and *s_M_*, respectively), of which a fraction of the CD4+ T cells (*f_T_*) and macrophages (*f_M_*) exiting from the bone marrow are CCR5-deficient, while the remainder are not.

### Key metrics of viral rebound

We defined *viral rebound* as occurring if the viral load surpasses the limit of detection of <50 copies of viral RNA/mL following the cessation of cART. Virtual patients who do not exhibit rebound after cART cessation and throughout the remainder of the simulation are assumed to be in remission and classified as responding positively to HSCT treatment. Time to viral rebound is measured as the duration from cART cessation to the onset of viral rebound for a given patient.

### Analysis of parameter importance

We use multiple methods to assess the relevance of parameters to the outcomes of the model and their correlations to each other. The Pearson correlation coefficient measures whether and to what extent two variables are linearly dependent and ranges between -1 and +1, with -1 indicating perfect inverse relationship, 0 indicating no relationship, and +1 indicating perfect direct relationship. For local sensitivity analysis, we use relative sensitivity defined as the percentage change in a key metric with respect to a percentage local perturbation in one parameter value. We use a parameter value change of 5% for these simulations. Similar to the correlation coefficient, a negative sensitivity value indicates an inverse relationship, 0 indicates no relationship, and a positive sensitivity value indicates a direct relationship. For instance, a relative sensitivity of 1 means that the 5% local perturbation in the particular model parameter results in a 5% increase in the selected key metric. For global sensitivity analysis, we use the Sobol index, a variance-based method. Each Sobol index represents the fraction of variance in the key metric contributed by the variance of a model parameter. First-order Sobol index measures solely for the effect of the model parameter alone while total Sobol index accounts for the interactions between model parameters on top of the first-order effect.

### Partial least squares discriminant analysis (PLS-DA)

Partial least squares discriminant analysis is a variant of the partial least squares (PLS) regression adapted to perform classification tasks, where the response variable is categorical. Similar to PLS regression, PLS-DA constructs latent variables that are linear combinations of the parameters to maximize the covariance between parameter variables (model parameters) and the categorical response variable (viral rebound vs. non-rebound). Here, PLS-DA allows us to project patient model parameters onto a lower-dimensional space spanned by the latent variables while maximizing the separation between the patients experiencing viral rebounding and those who do not. The degree of separation in the score plot between the two subpopulations indicates how well the variability in the response variable can be explained by the variability in the parameter variables. Concurrently, the loading plot illustrates the direction and extent to which a given parameter variable drives the separation between the two subpopulations.

## Results

### cART, cART cessation, and viral rebound

To illustrate the effects of combination antiretroviral therapy (cART) alone on infection trajectory, we first performed simulations using the mean parameter values and initial conditions across all patients (Figure 2). We simulated three different scenarios: infection only with no treatment, infection followed by indefinite cART that begins on day 200, and infection followed by 500 days of cART starting from day 200. Without treatment, HIV infection typically leads to a sudden surge in viral load and a decline in CD4+ T cell count. In our simulations, cART is effective in reducing the viral load; not eliminating it, but maintaining it at a low level, generally below the level of detection (Figure 2A), and CD4+ T cell counts recover (Figure 2B). However, following treatment interruption (TI), the termination of cART leads to rapid rebound of viral load back to the pre-cART levels and a decline in CD4+ T cells. This matches real-world experience with rebound after TI ^12^. To understand interindividual variability (IIV) in the effects of cART, we simulated similar treatment across a large population of virtual patients, each virtual patient having individual parameter values and initial conditions. The virtual patients were divided into four separate groups based on the time from HIV seroconversion to AIDS diagnosis: less than 3.5 years (rapid progressors, Population 1), between 3.5 and 7 years (Population 2), between 7 and 9 years (Population 3), and more than 9 years (slow progressors, Population 4). When cART is given from days 200 to 1300 post-infection, individual simulations revealed similar trajectories as the average simulation in that cART significantly reduces the viral load while interruption of cART leads to viral rebound (Figure 3A). To show that this pattern does not depend on the rate of AIDS progression or the duration of cART, we performed individual simulations on all four virtual populations and tested different durations of cART from 3 to 50 years. Across the virtual populations, we found that termination of cART consistently leads to viral rebound regardless of the length of cART treatment (Figure S1). Increasing the period of cART treatment does marginally increase the time to viral rebound; the variance in time to rebound also increases with cART duration (Figure 3b). But ultimately, cessation of cART results in viral rebound for all, even after 50 years of cART.

**Figure 2.**
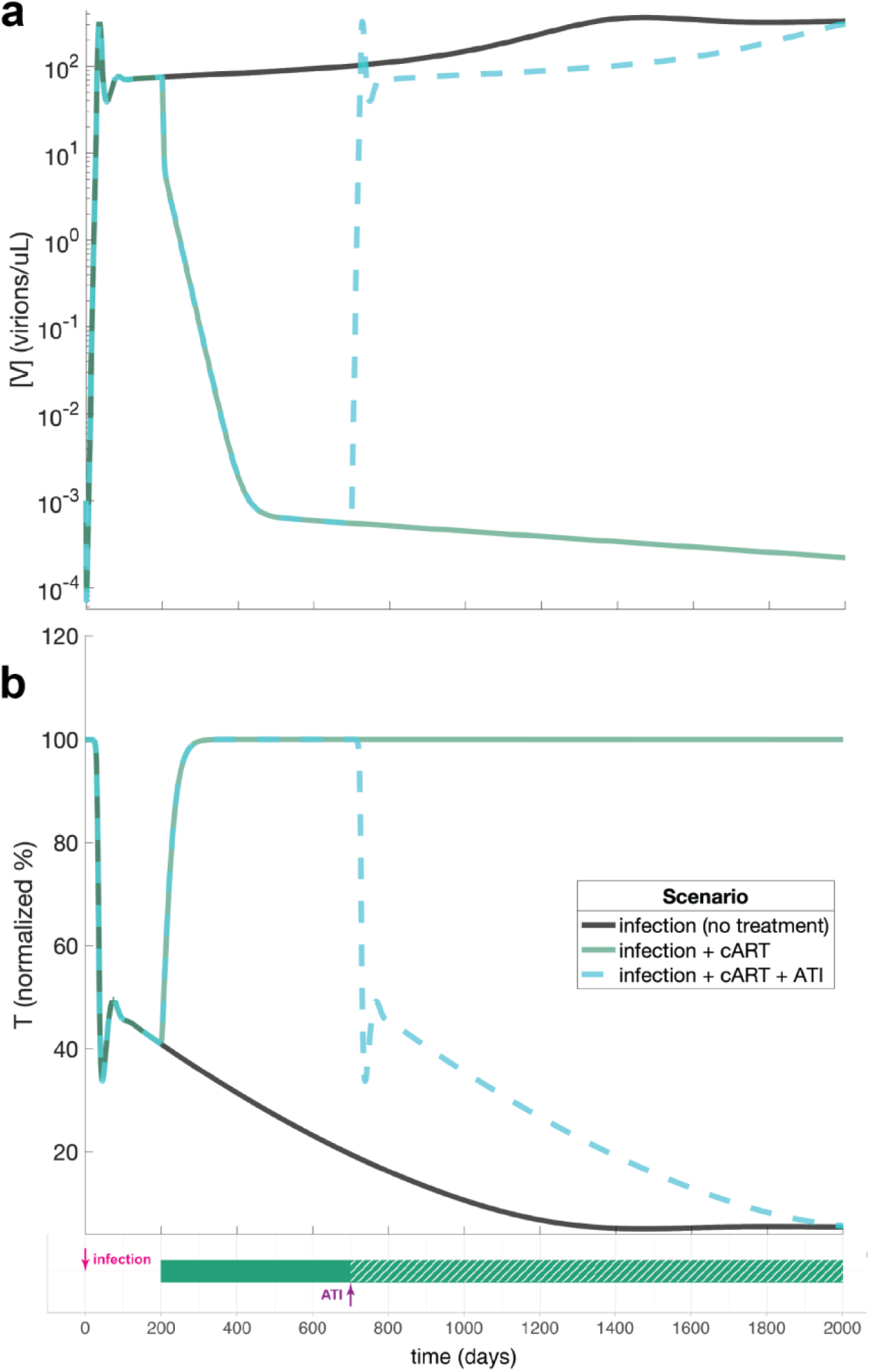
Simulations of typical time courses of HIV infection, cART, and treatment interruption (TI). (**a**) Predicted viral load over time following infection and cART treatment. (**b**) CD4+ T cell counts over time following infection and cART treatment. CD4+ T cell counts were normalized by the number of initial CD4+ T cells present in the system. Parameters used for this simulation are the mean values across the four virtual populations. Three different scenarios are simulated: infection only (no treatment); infection followed by indefinite cART; and infection followed by cART that ends (treatment interruption, TI). In these simulations, infection occurs on day 0, cART begins on day 200, and treatment interruption occurs on day 700.

**Figure 3.**
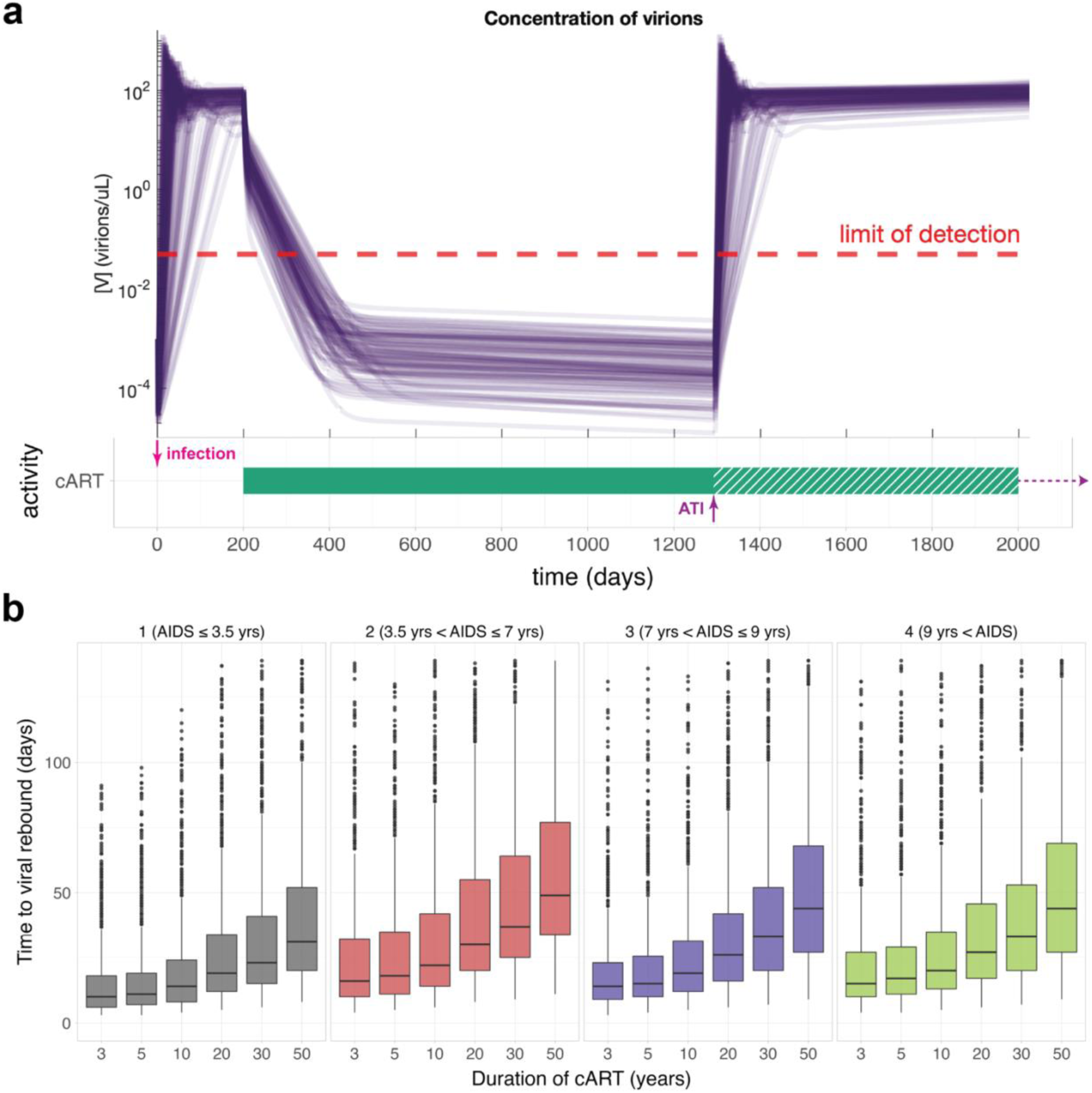
Simulations of infection, cART, and treatment interruption (TI) in the virtual patient population. (**a**) Predicted viral loads over time for each of 100 virtual patients in Population 1 (rapid progressors). The virtual clinical trial consists of: infection at day 0; cART from day 200; and TI (cessation of cART) at day 1300. (**b**) Population distributions of time to viral rebound (T_VR_) for different durations of cART. The threshold for viral rebound is defined by the limit of detection at <50 copies/mL. Time to viral rebound measures the length of time it takes for each patient to exceed the limit of detection starting from TI. The boxes represent the 25^th^ to 75^th^ percentiles, and the bolded line marks the median of the patient distribution. Outliers are shown in black dots; the edge of the whiskers mark the minimum and maximum of the patients considered.

### Biomarkers of time to rebound following cART cessation

To examine how the variance of cART treatment outcome is associated with the IIV among the virtual patients, we calculated population-wise Pearson correlation coefficients between key metrics and model parameters (Figure 4). Three key metrics were selected to characterize each infection trajectory: viral load immediately prior to TI ([V]_pre-TI_), viral load at the end of simulation ([V]_post-TI_), and time to viral rebound (T_VR_). We found that most model parameters are weakly correlated with the chosen key metrics in most virtual populations, with the exception of *k_L_* in Population 1. *k_L_* determines the fraction of viral infection that leads to viral latency and shows strong positive relationship with the [V]_pre-ATI_ in Population 1. This could be explained by the importance of the latent viral reservoir in sustaining the viral load as cART actively targets new infections but does not target cells that are already infected. To go one step further, we explored whether the behavior of the infection as a system is responsive to the changes in specific model parameters. We conducted population-wise local sensitivity analysis of the aforementioned key metrics with respect to the model parameters. Unlike the correlation coefficient, the relationship measured by sensitivity analysis need not be linear. By looking at the population mean local sensitivities, we found that most influential parameters primarily fall into two groups: increase in the parameters leads to increase in pre-ATI and post-ATI viral loads and decrease in time to viral rebound, and vice versa, increase in the parameters leads to decrease in pre-ATI and post-ATI viral loads but increase in time to viral rebound (Figure 5). A third group consists of parameters that sparsely impact one particular key metric but not the other two. A successful therapeutic outcome is one where pre-ATI and post-ATI viral loads are maintained at a low level while time to viral rebound is lengthened. Parameters that, when increased in value, increase viral loads and decrease time to viral rebound (both unfavorable for the patient) include the infection rate of virus on uninfected CD4+ T cells (*k_VT_*) and the burst size of infected CD4+ T cells (number of viruses produced by infected CD4+ T cells, *n_T_*). An increased *k_VT_* leads to more CD4+ T cells being productively infected while an increased *n_T_* leads to more progeny virions being generated from these productively infected cells; both of these processes directly increase the viral load, thus shortening the time to viral rebound. On the contrary, parameters that, when increased, decrease viral loads and increase time to viral rebound include the rates of death of uninfected CD4+ T cells, infected CD4+ T cells, and viruses (*d_T_*, *d_TI_*, and *d_V_* respectively). An increased *d_T_* results in fewer target cells available for the viruses to infect while an increased *d_TI_* results in fewer progeny virions produced by infected cells. Along with an increase in death rate of viruses, all three processes directly result in a diminished viral load, thus extending the time to viral rebound. The third group includes parameters such as the maximum increase in proliferation rate of uninfected macrophages under inflammatory response to infection (*p_M_*), the burst size of infected macrophages (*n_M_*), the rates of death of uninfected and infected macrophages (*d_M_* and *d_MI_* respectively), rate of activation of latently infected CD4+ T cells (*a_L_*), and the fraction of latent infections from infected CD4+ T cells (*k_L_*). Increase in *a_L_* and *k_L_* results in an expansion of the latent viral reservoir; the latent viral reservoir evades suppression under cART and becomes an important scaffold for viral rebound once treatment is ceased. *p_M_*, *n_M_*, *d_M_*, and *d_MI_* all impact the level of macrophages, which play an important role as secondary targets for new infections once the viruses have exhausted the uninfected CD4+ T cells; this is particularly the case in the absence of treatment. Similar to the variance in cART treatment outcomes, we also observed patient heterogeneity in the population distribution of local sensitivities among different parameter-metric combinations (Figure S2). Given that there are multiple parameters to which the system displays strong local sensitivity, why then do we see only weak correlations between these same parameters and the output metrics across the population? Most likely, the difference is that the sensitivity is high when varying the parameters one-at-a-time, but between virtual patients, multiple of these parameters are different simultaneously, canceling each other out or each contributing part and none contributing all of the differences in outcomes between the patients.

**Figure 4.**
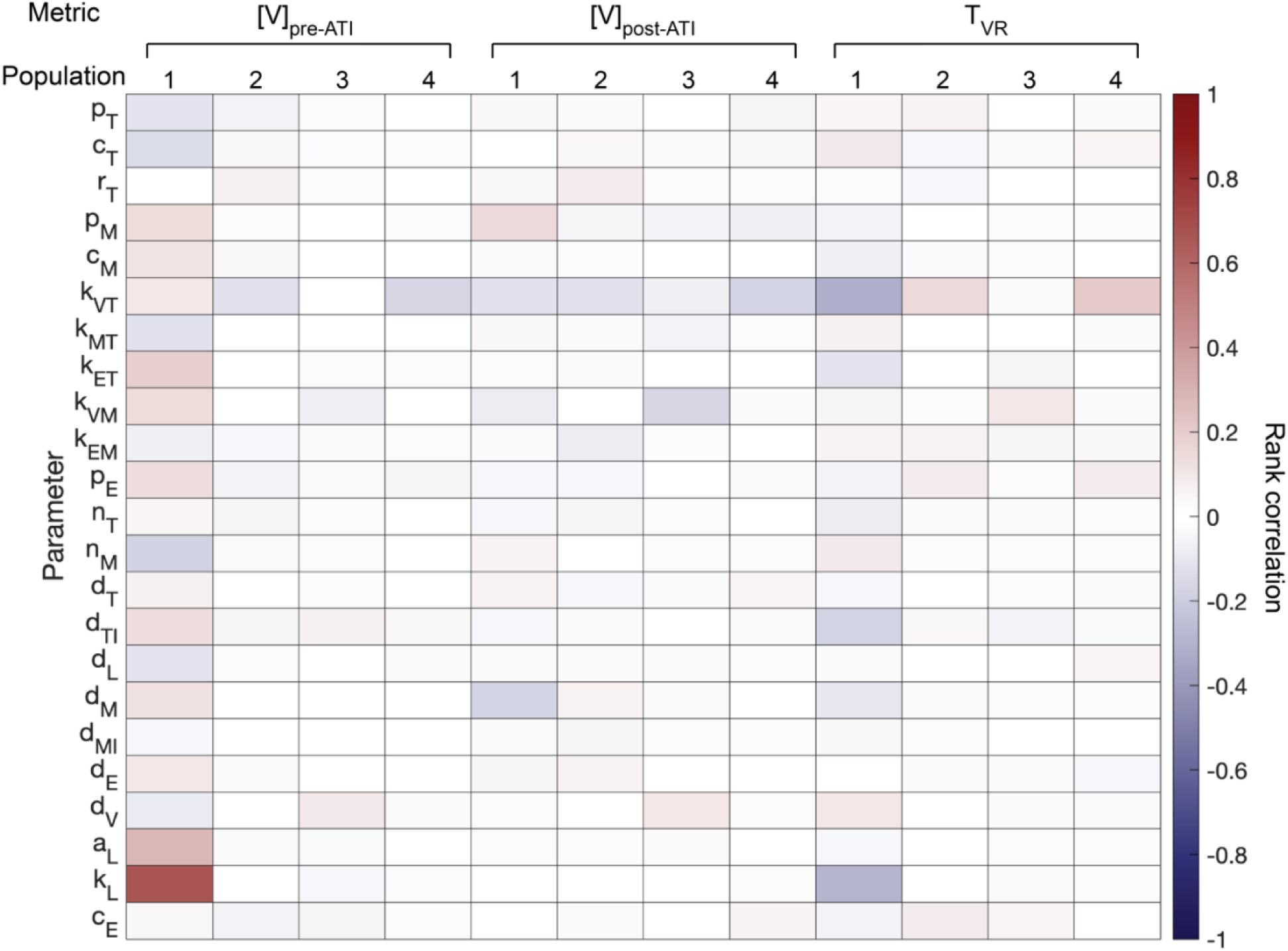
Population-wise correlation between key output metrics (viral load and time to rebound) and model parameters. Pearson correlation coefficient measures the extent of linear dependence between two random variables; negative values (blue) indicate inverse relationship, positive values (red) indicate direct relationship, and 0 (white) indicates no relationship. Each cell of the heatmap represents the correlation between one parameter and one key metric in one of the four virtual populations. Three key metrics are evaluated for each virtual patient: viral load immediately prior to ATI ([V]_pre-ATI_), viral load at day 2000 ([V]_post-ATI_), and time to viral rebound (T_VR_).

**Figure 5.**
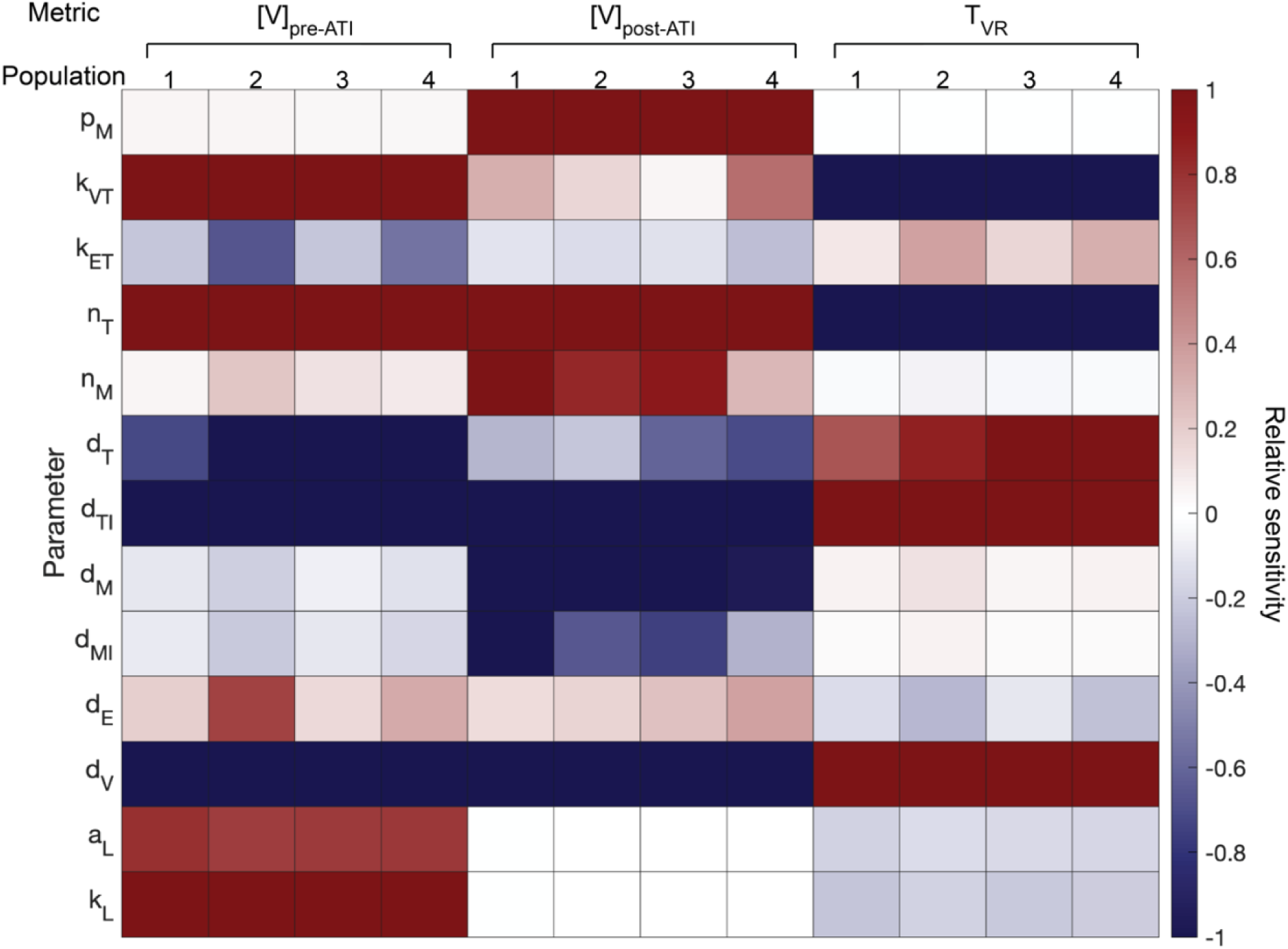
Population-wide local sensitivity analysis of key metrics on model parameters. Relative sensitivity is defined as the percent change in key metric following 5% increase in a parameter value. Negative sensitivity values (blue) indicate inverse relationship, positive sensitivity values (red) indicate direct relationship, and 0 (white) indicates no relationship. Each cell of the heatmap represents the mean relative sensitivity of one parameter with respect to one key metric across each of the four virtual populations. The virtual clinical trial consists of: infection at day 0; cART from day 200; and TI (cART cessation) at day 1300. Only parameters with the top one-third sensitivity values are shown. Three key metrics are evaluated for each virtual patient: viral load immediately prior to ATI ([V]_pre-ATI_), viral load at day 2000 ([V]_post-ATI_), and time to viral rebound (T_VR_).

To determine how much the contribution in key metric variances is due to the parameter alone or through interactions with other parameters, we conducted a global sensitivity analysis by calculating the first order and total Sobol indices for the same parameter-metric combinations (Figure 6). We evaluated the first order and total Sobol indices jointly on patients from all four virtual populations (Combined Population). Out of the three key metrics, we found *k_L_* (the fraction of viral infection leading to viral latency) to be most influential on pre-ATI viral loads, *p_M_* (the proliferation rate of uninfected macrophages under inflammation) to be most influential on post-ATI viral loads, and *k_VT_* (the infection rate of virus on uninfected CD4+ T cells) to be most influential on time to viral rebound based on first order effect alone. However, all of the influential parameters have a total Sobol index that significantly outweighs the corresponding first order Sobol index. Noticeably, the total effect of the infection rate of virus on uninfected macrophages (*k_VM_*) far surpasses that of *p_M_* as the top contributor for post-ATI viral loads. As previously mentioned, *k_L_* is important in replenishing the latent viral reservoir while escaping the suppression by cART. *k_VT_* plays a role in determining the number of infected CD4+ T cells in the system, which directly impacts the viral loads and time to viral rebound. *k_VM_* and *p_M_* regulate the availability of macrophages as secondary targets for new infections, especially once the viruses have exhausted the uninfected CD4+ T cells.

**Figure 6.**
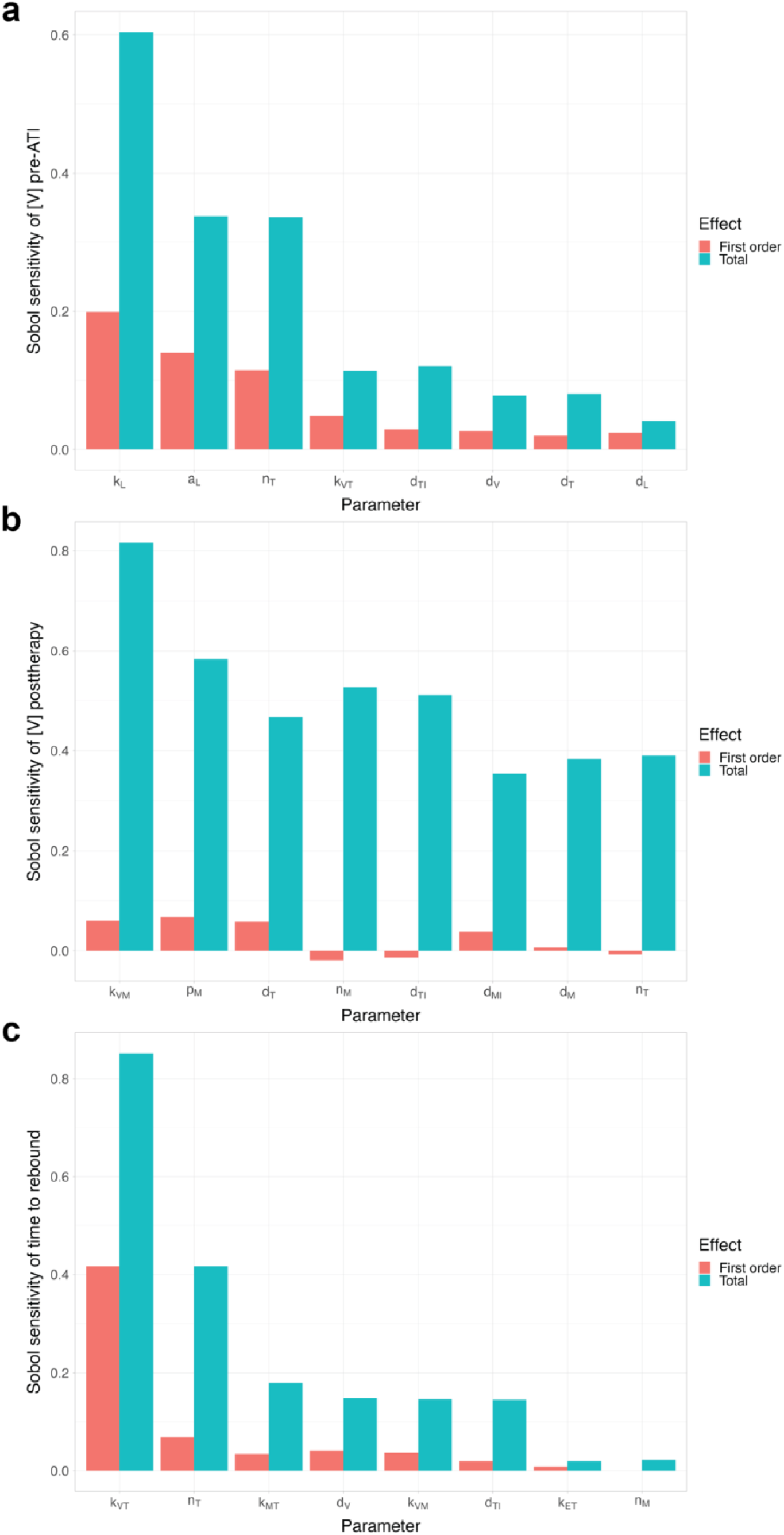
Global sensitivity analysis of key metrics to model parameters in the combined virtual population. Sobol index is a variance-based measure of global sensitivity: the fraction of variance in each metric that is explained by variance of each model parameter. First-order Sobol index accounts solely for the effect of one model parameter, while total Sobol index considers the first-order effect along with all interactions with other model parameters. The virtual clinical trial consists of: infection at day 0; cART from day 200; and TI (cART cessation) at day 1300. Only parameters with top one-third sensitivity values are shown. Three key metrics are evaluated for each virtual patient: viral load immediately prior to ATI ([V]_pre-ATI_), viral load at day 2000 ([V]_post-ATI_), and time to viral rebound (T_VR_).

To better illustrate how these key parameters impact on the output metrics, and on the infection trajectories generally, we visualized the simulated viral loads for 100 patients in each of the virtual populations as part of a virtual clinical trial *in silico* (Figure 7A,C,E; Figures S3-5, left panels). The virtual clinical trial consists of three sequential periods: (1) untreated infection starting at day 0; (2) treatment with cART starting at day 200; and (3) return to no treatment (i.e. cessation of cART) starting at day 1300. To visualize the contributions of parameters on the key metrics, we colored the individual patient trajectories based on the rank (across the population) of the previously identified top contributing parameters from the global sensitivity analysis to emphasize different features of the simulations: *k_L_* for pre-ATI viral loads, *k_VM_* for post-ATI viral loads, and *k_VT_* for time to viral rebound. The three key metrics were segmented out and enlarged into inset reference windows to highlight the parameter gradients across the patient population (Figure 7A, C, E, inset rectangles). A highly predictive parameter-metric relationship would appear as a monotonic gradient in our simulations. The simulations revealed partial (noisy) gradients for the linkage between *k_L_* and pre-ATI viral loads as well as the linkage between *k_VT_* and time to viral rebound. In contrast, the window for *k_VM_* and post-ATI viral loads revealed minimal gradients. Concurrently, we also plotted the key metrics directly against the suggested driving model parameters (Figures 7, S3-5, right panel). Most direct comparisons showed a weak linear relationship between the key metric and the driving parameter with the exception of *k_L_* and pre-ATI viral loads in Population 1.

**Figure 7.**
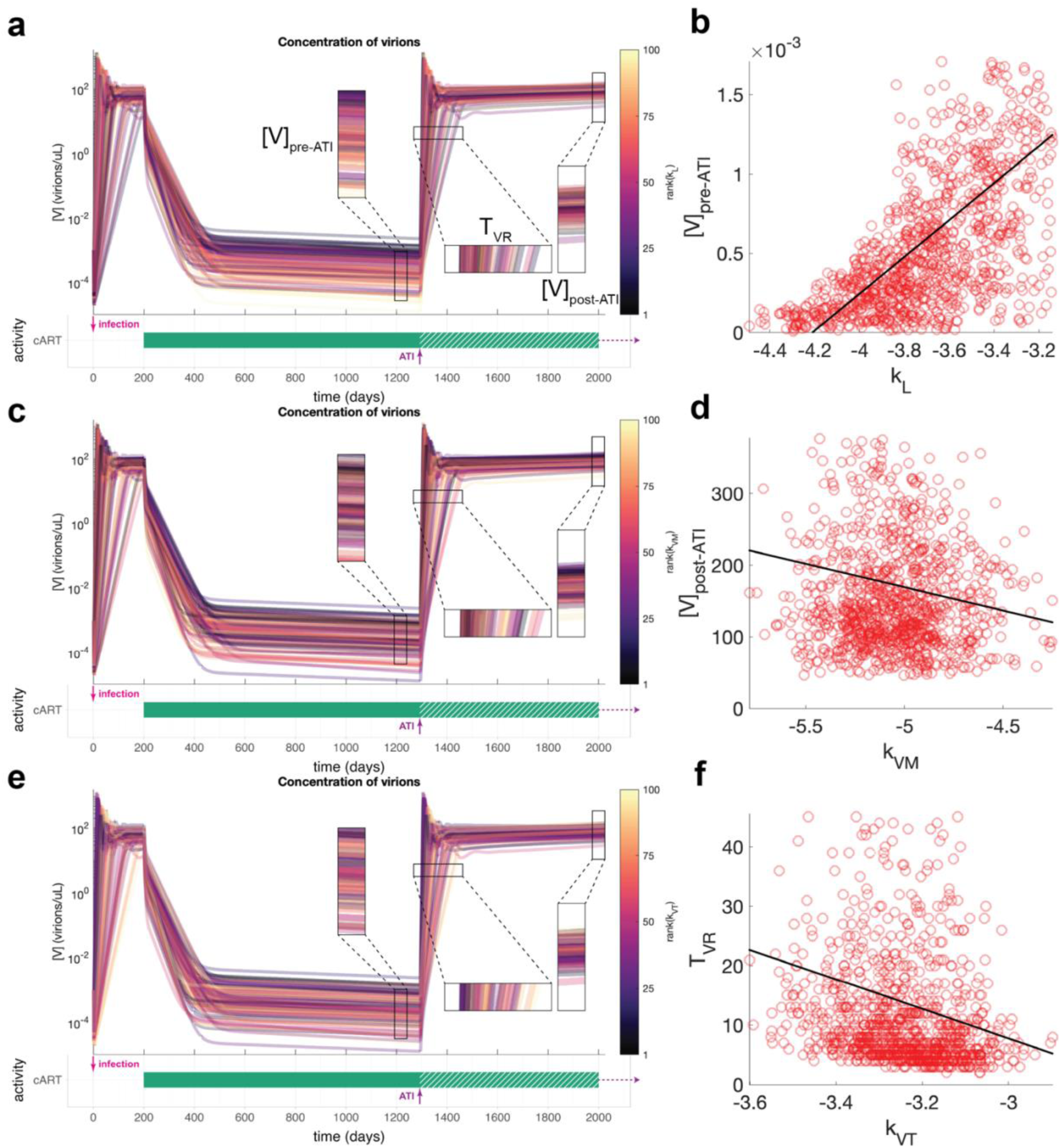
Simulations of infection trajectories across the virtual population. (**a,c,e**) Viral loads for 100 virtual patients in Population 1. The virtual clinical trial consists of: infection at day 0; cART from day 200; and TI (cART cessation) at day 1300. Individual patient trajectories are colored by the rank of different model parameters to emphasize distinct features of the simulations: (**a,b**) rate of CD4+ T cell latency conversion (*k_L_*) for viral load immediately prior to ATI ([V]_pre-ATI_); (**c,d**) rate of macrophage infection by virus (*k_VM_*) for viral load at day X post-TI ([V]_post-ATI_); (**e.f**) and rate of CD4+ T cell infection by virus for time to viral rebound (*T_VR_*). These were the top driving model parameters identified for each key metric in the Sobol global sensitivity analysis. (**b,d,f**) Direct comparison of key metrics with suggested driving model parameters. Each red circle is one virtual patient, and the line of best fit is shown in black.

In the prior analyses, we have assessed the parameter-metric relationships while singling out one model parameter and one key metric for each iteration. To determine the independence among the model parameters and among the key metrics, we computed within-group Pearson correlation coefficients for the parameters and the metrics (Figure 8). We found that most parameters are weakly correlated with the others with the exceptions of burst size (*n_T_*) being inversely correlated with rate constant of viral infection of T cells (*k_VT_*), and the rate of death of CD8+ T cells (*d_E_*) being directly correlated with the proliferation rate of uninfected CD4+ T cells under homeostasis (*r_T_*). The inverse relationship between *n_T_* and *k_VT_* may be an artifact from the parameter optimization where the values of these two parameters are jointly constrained by the levels of viral replication observed in patients. The ratio of CD4+ T cells to CD8+ T cells has been established as an indicator for the strength of one’s immune function ^13^. A CD4+/CD8+ ratio of less than 1 suggests that the individual’s immune system has been compromised as in the case of HIV patients. The positive relationship between *d_E_* and *r_T_* could be thought of as a restorative mechanism to maintain equilibrium between different subsets of T cells under homeostatic conditions ^14^. Among the key metrics, we found pre-ATI and post-ATI viral loads to be slightly positively correlated with each other. Both the pre-ATI and post-ATI viral loads were found to be inversely correlated with time to viral rebound; this was expected as higher viral loads exerts a higher burden on the immune system, leading to faster viral rebound.

**Figure 8.**
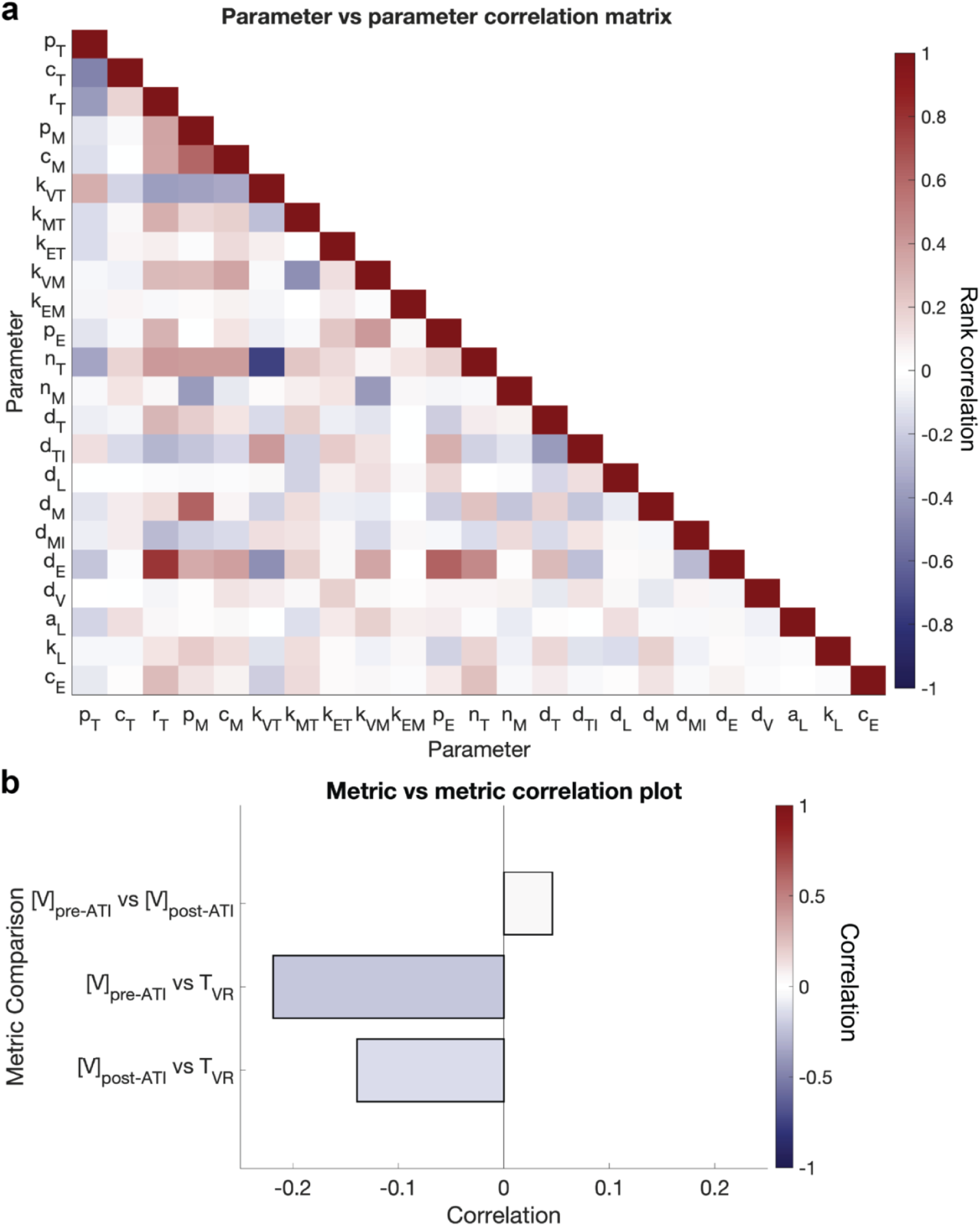
Correlations among model parameters and among key metrics across the combined virtual populations. (**a**) Correlations among model parameters. (**b**) Correlations between key metrics (virtual clinical observations). Pearson correlation coefficient is described in Figure 4.

### Biomarkers of response to augmented HSCT

Given that cessation of cART inevitably leads to viral rebound, we hematopoietic stem cell transplantation (HSCT) as a potential cure for HIV infection. To date, this combination has been proven successful in achieving HIV remission in five clinical patients. The transplantation procedure, results in the recipient having a chimeric immune system consisting of both wild-type (host-derived) and augmented (donor-derived) immune cells. To study the role that the chimeric immune system plays in determining the therapeutic outcome, we simulated different levels of chimerism using individual parameter sets and initial conditions from Population 1 under a virtual clinical trial treatment scenario: infection followed by 500 days of cART starting from day 200, engraftment of HSCT starting at day 500, followed by cART cessation at 700. To represent different levels of chimerism, we varied the percentage of augmented immune cells present in the chimeric immune system, from 25% to 100%. We found that the proportion of virtual patients experiencing viral rebound decreases as the fraction of augmented cells increases in the chimeric immune system (Figure 9a). When the chimeric immune system is entirely composed of augmented immune cells, our simulations predicted that all of the patients will achieve HIV remission. We were able to replicate this finding in the other three virtual populations; moreover, other than the rapid progressors in Population 1, patients from Populations 2-4 are predicted to achieve remission even when the augmented cells constitute only up to 75% of the chimeric immune system (Figure 9b). We then tested how the treatment outcomes of HSCT are affected by different durations of cART. Hence, we performed simulations as described above for individuals undergoing HSCT, assuming half of the chimeric immune system is composed of augmented immune cells, and tested durations of cART from 3 to 50 years. We observed no significant change in percentage of patients predicted to experience viral rebound as we prolonged the duration of cART (Figure 10a). This is consistent across all four virtual populations regardless of the rate of AIDS progression. Similar to the simulations without cART, increasing the duration of cART increases (though only slightly) the median time to viral rebound as well as its variance, in that fraction of patients predicted to rebound (Figure 10b). One slight exception to this effect seems to be within Population 4, the slowest progressors, where both the highest median time to viral rebound and variance are reached when patients are treated with cART for 20 years.

**Figure 9.**
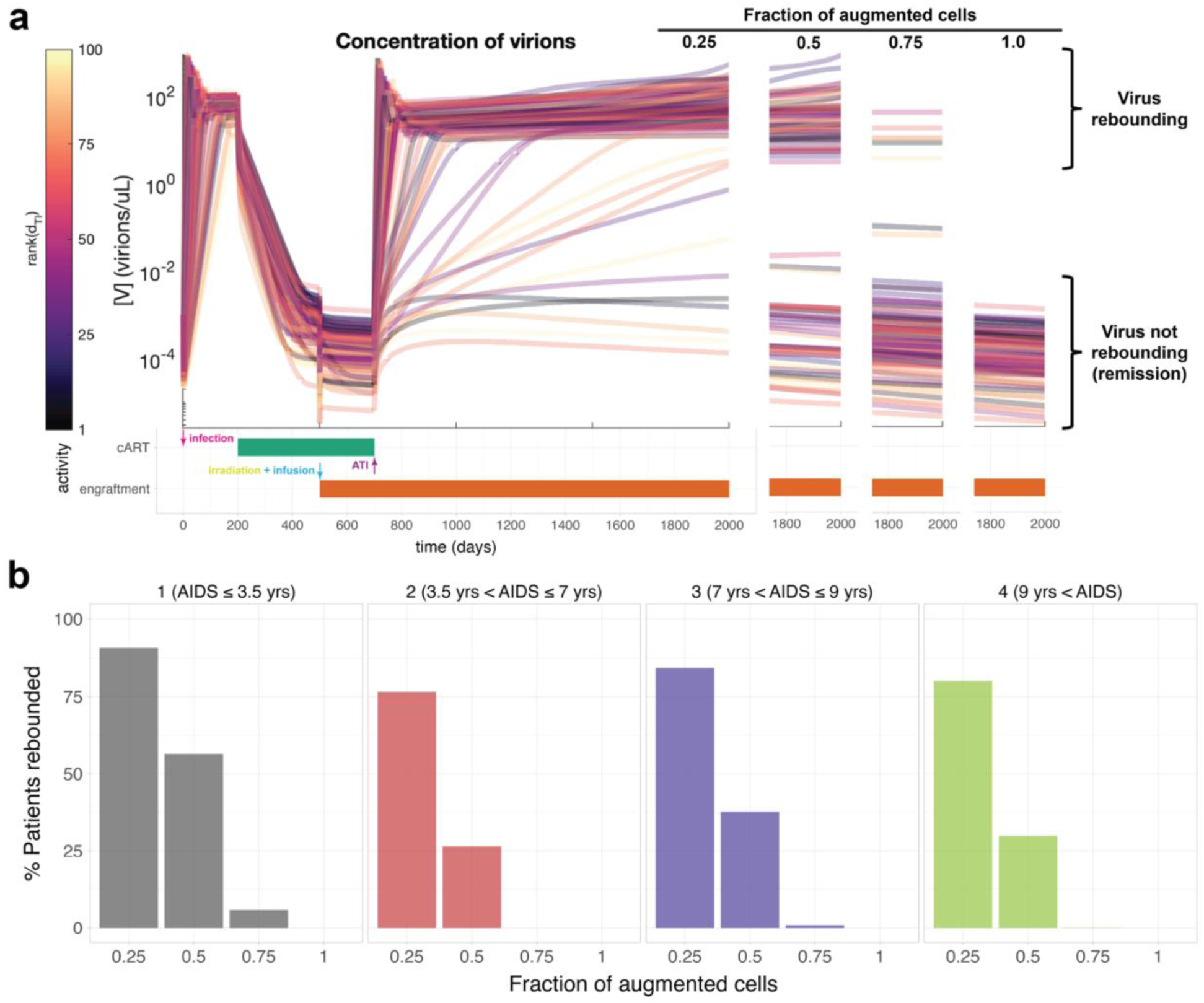
Effects of chimerism on hematopoietic stem cell transplantation (HSCT) as a treatment for HIV-infected patients. (**a**) Simulated viral loads for 100 virtual patients in Population 1. Individual patient trajectories are colored by the rank (across the virtual population) of the death rate of infected CD4+ T cells (*d_TI_*). The virtual clinical trial consists of: infection at day 0; cART from day 200; HSCT at day 500; and TI (cART cessation) at day 700. Four chimerism scenarios are simulated (augmented cells:wild-type cells): 25%:75%; 50%:50%; 75%:25%; and 100%:0%. The virtual population is stratified into rebounders and non-rebounders (remission) based on the viral loads at day X post-TI. The threshold for viral rebound is defined by the limit of detection at <50 copies/mL. (**b**) Percent of patients in each virtual population predicted to experience viral rebound under different levels of chimerism post-HSCT.

**Figure 10.**
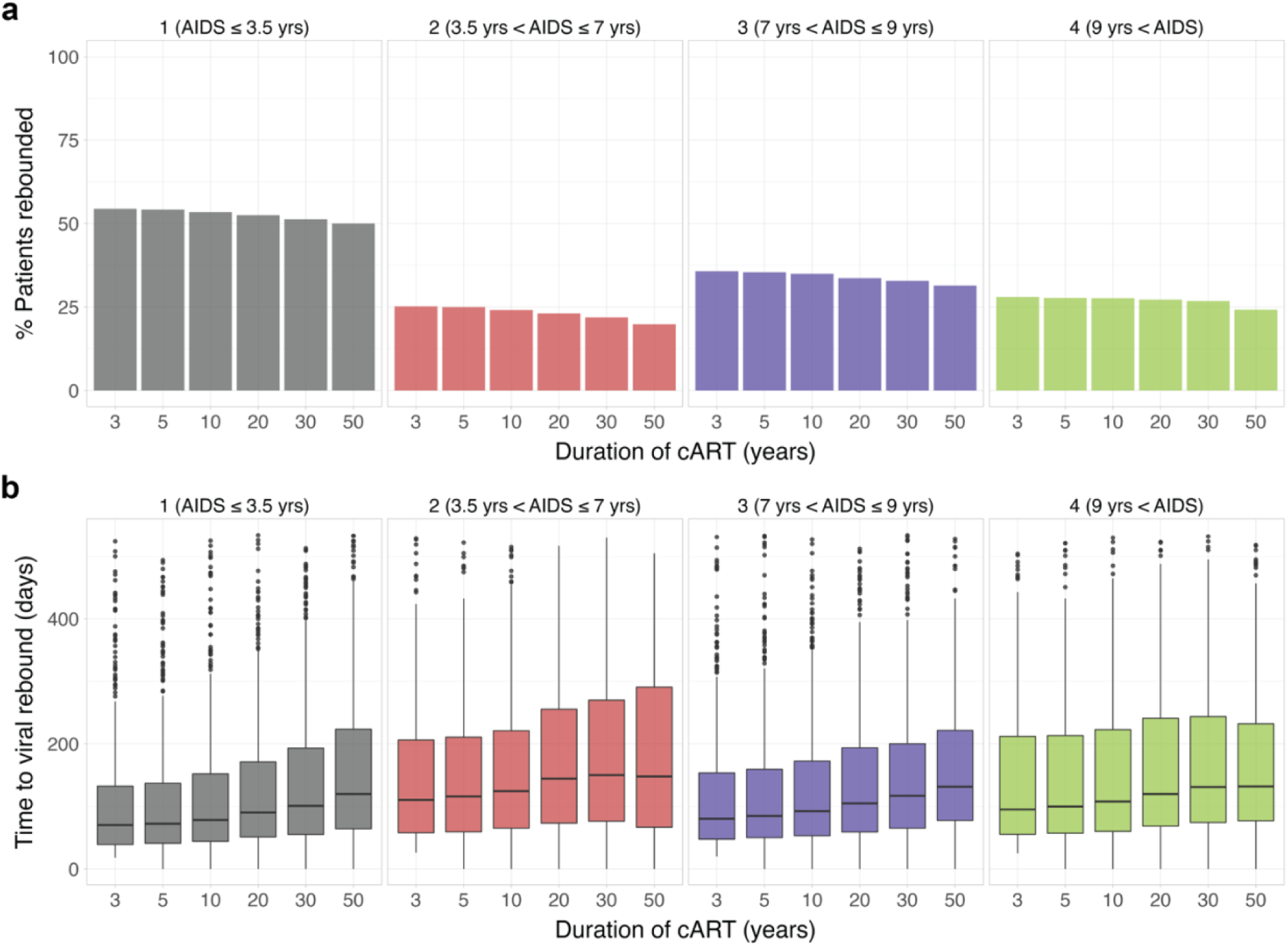
Effects of extending cART duration on hematopoietic stem cell transplantation (HSCT) as an HIV treatment. (**a**) Percent of patients in each virtual population experiencing viral rebound following cessation of cART, following HSCT and treatment with cART for different lengths of time (3-50 years). The virtual clinical trial consists of: infection at day 0; cART from day 200; HSCT at day 500; and TI (cART cessation) at a day corresponding to the specified cART duration. The chimeric immune system is assumed to include 50% augmented cells and 50% wild-type cells. The virtual populations are stratified into rebounders and non-rebounders (remission) based on the final viral loads. The threshold for viral rebound is defined by the limit of detection at <50 copies/mL. (**b**) Boxplots depict the population distributions of time to viral rebound under each of the scenarios. Time to viral rebound measures the length of time it takes for the patient to exceed the limit of detection starting from TI. The box plot represents the 25th percentile, median, and 75^th^ percentile of the distributions.

Similar to the simulations under cART treatment alone, we explored how model parameters drive the population variance in the therapeutic outcomes under the combination of HSCT and cART. Assuming 50% augmented immune cells in the chimeric immune system, we observed a bimodal response among the patients, where a fraction of the virtual population is predicted to experience viral rebound while the remainder is not. Hence, unlike the key metrics selected previously for cART alone, patient viral rebound shows a measurable binary response. To ascertain how well model parameters explain this variability in patient viral rebounding, we performed Partial Least-Squares Discriminant Analysis (PLS-DA) using model parameters as features and patient viral rebounding as a binary label. By projecting the patients onto a lower dimensional space spanned by the latent variables, we found that the patient rebounding and non-rebounding subpopulations are not easily separable based on linear combinations of the model parameters (Figures 12, S6-S10, right two columns). Scree plots in the parameter space confirmed our previous finding from inter-parameter correlation coefficients that there is minimal redundancy in the information carried by each model parameter (Figures S6-S10, bottom row). Moreover, scree plots in the response space showed similar results as the prior global sensitivity analysis in that there is not one model parameter that single-handedly influences the infection trajectories; rather, infection trajectories are determined primarily by the interactions between model parameters (Figures S6-S10, bottom row). This is validated by the regression coefficient plot, which presents most of the model parameters having only small individual impacts on the direct predictions of viral rebounding responses (Figure 11a).

**Figure 11.**
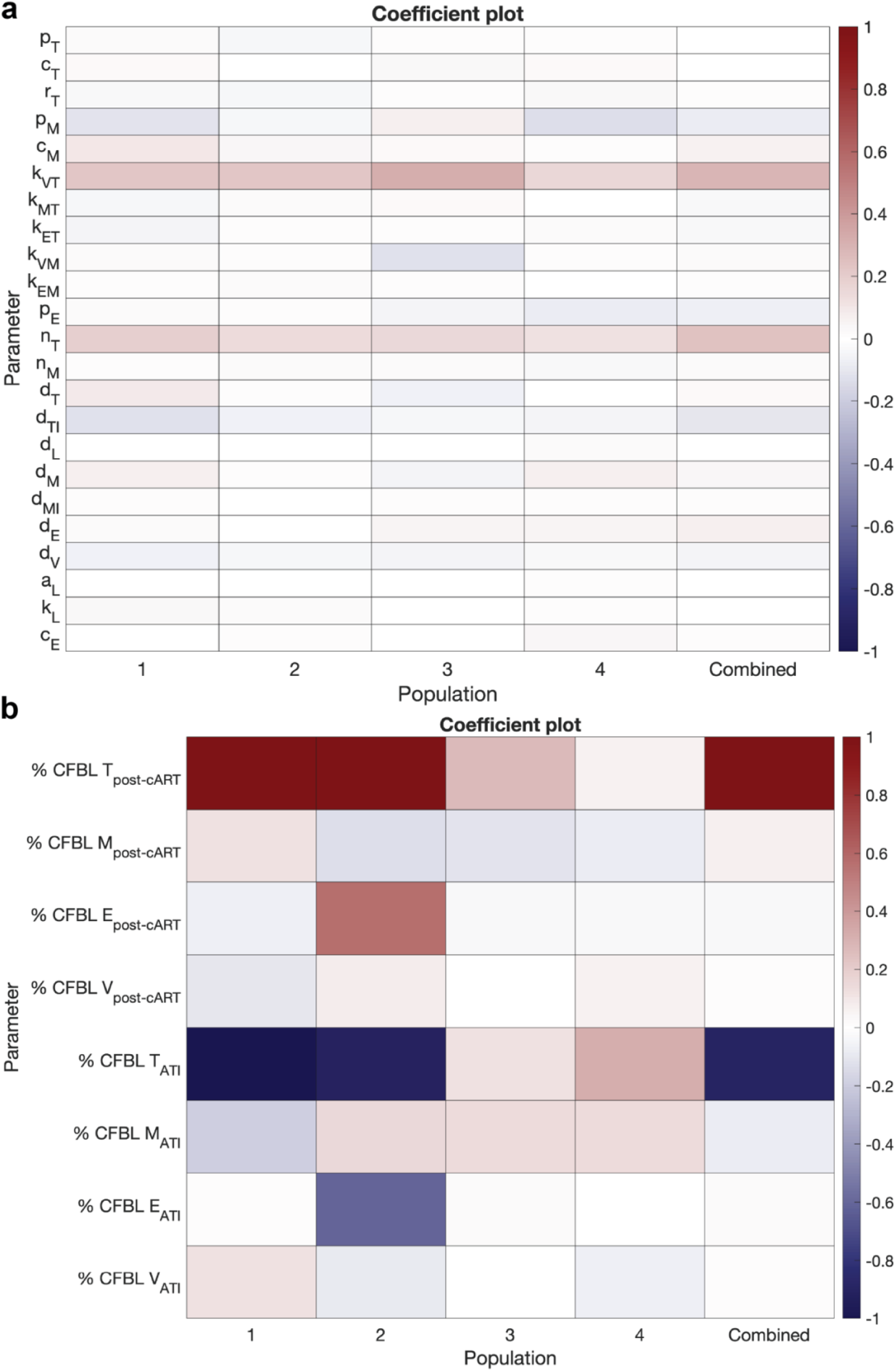
Coefficients of model parameters and key output metrics (viral load and time to rebound) in PLS-DA regression models. (**a**) Coefficients of mechanistic model parameters as drivers of therapeutic success of HIV-infected patients undergoing hematopoietic stem cell transplantation (HSCT) using partial least squares (PLS) regression. (**b**) Coefficients of virtual clinical observations for PLS. Coefficient indicate how each input variables is related to the response variable (viral rebound vs remission). Negative coefficients (blue) indicate inverse relationship, positive coefficients (red) indicate direct relationship, and 0 (white) indicates no relationship. The four virtual populations and the combined population are shown separately.

**Figure 12.**
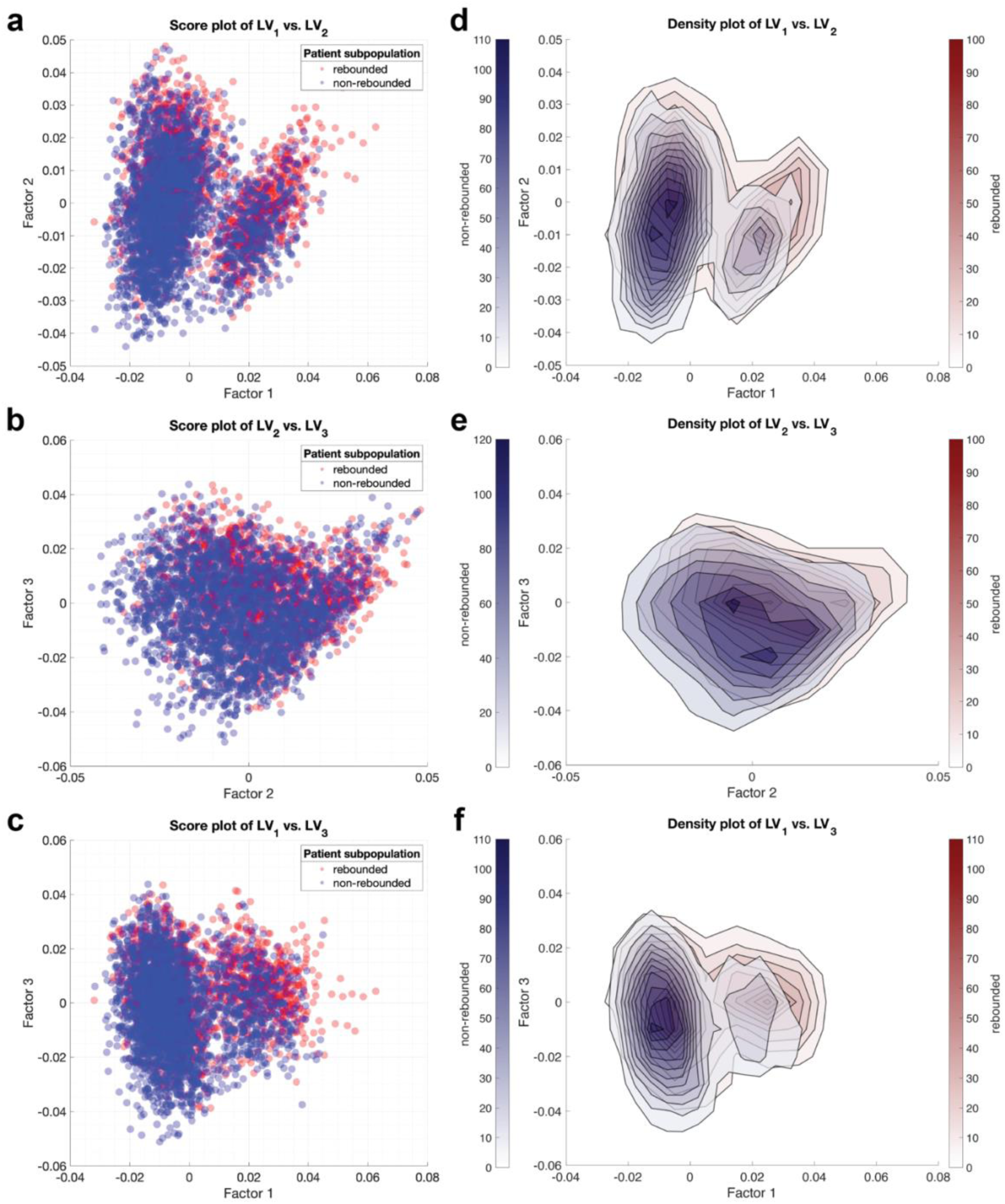
Model parameters as drivers of HSCT therapeutic success of HIV-infected patients in the combined population. The virtual patients undergoing hematopoietic stem cell transplantation (HSCT) were segregated into responders (no viral rebound, blue) and nonreseponders (viral rebound, red) based on the final viral loads. The threshold for viral rebound is defined by the limit of detection at <50 copies/mL. Partial least squares (PLS) regression used to identify latent variables that best separate the groups. (**a,b,c**) Score plot (one dot = one virtual patient) with respect to two of the first three latent variables. (**d,e,f**) Density plots corresponding to the patient scatter plots shown in the left column.

Although we were unable to pinpoint a singular driving force among the model parameters, the strong correlations between the key metrics as shown previously suggested that virtual clinical observations could be plausible determinants of population variability in therapeutic outcomes. To draw a distinction between model parameters and virtual clinical observations, model parameters are inputs to our model that defines the infection dynamics within a patient, whereas virtual clinical observations, or previously key metrics, are outputs of simulations based on our model. In practice, these virtual clinical observations are quantities that are more accessible and could be available to be measured clinically in each patient. Twelve clinical observations were defined based on the model outputs at various critical time points of the treatment scenario: total CD4+ T cell counts, macrophages counts, CD8+ T cell counts, and viral loads immediately prior to cART ([T]_pre-cART_, [M]_pre-cART_, [E]_pre-cART_, and [V]_pre-cART_ respectively), after cART ([T]_post-cART_, [M]_post-cART_, [E]_post-cART_, and [V]_post-cART_ respectively), and at the end of simulation ([T]_TI_, [M]_TI_, [E]_TI_, and [V]_TI_ respectively). We repeated PLS-DA on patient viral rebounding responses, but this time using these virtual clinical observations as features rather than the model parameters. Lower-dimensional projection of patients using latent variables consisting of linear combinations of clinical observations showed improved separations between the rebounding and non-rebounding patient clusters (Figures 13, S11-S15, right two columns). Scree plots in the parameter space reaffirmed our previous analysis on correlations between key metrics in that there exists some overlapping information among the clinical observations (S11-S15, bottom row). Moreover, scree plots in the response space are consistent with the lower-dimensional projections in score plots, indicating the patient rebounding and non-rebounding subpopulations are separable in two dimensions using latent variables derived from clinical observations (S11-S15, bottom row). The regression coefficient plot indicates that [T]_post-cART_ and [T]_ATI_ are strong predictors of the patient heterogeneity in viral rebound response (Figure 11b). Particularly, increase in [T]_post-cART_ drives viral rebounding while increase in [T]_ATI_ hinders viral rebounding. As mentioned previously, the maintenance of the latent viral reservoir plays a crucial role in providing a scaffold for viral rebound once cART treatment is terminated; the sustained presence of the latent viral reservoir correlates with an increase in [T]_post-cART_. On the other hand, an increase in [T]_TI_ suggests less depletion of CD4+ T cells and thus a lower likelihood of viral rebound.

**Figure 13.**
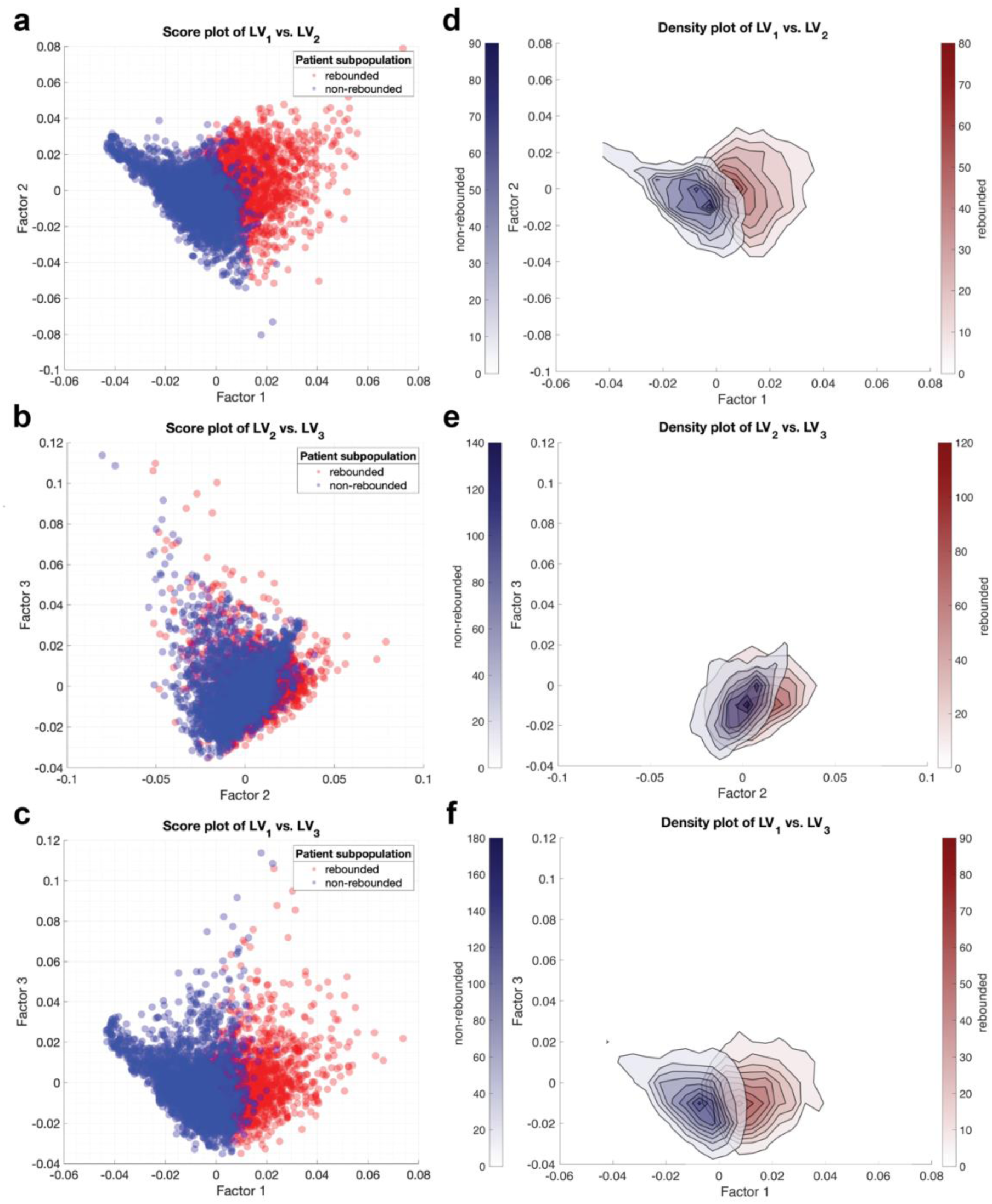
Clinical observations as drivers of therapeutic success of HIV-infected patients in the combined population. The virtual patients undergoing hematopoietic stem cell transplantation (HSCT) were segregated into responders (no viral rebound, blue) and nonreseponders (viral rebound, red) based on the final viral loads. The threshold for viral rebound is defined by the limit of detection at <50 copies/mL. Partial least squares (PLS) regression used to identify latent variables that best separate the groups. (**a,b,c**) Score plot (one dot = one virtual patient) with respect to two of the first three latent variables. (**d,e,f**) Density plots corresponding to the patient scatter plots shown in the left column.

## Discussion

In this study, we used a previously published mechanistic model of HIV dynamics, parameterized with virtual population parameters to represent the distribution of patient populations, to investigate the heterogeneity in viral rebound response following the cessation of cART,and the heterogeneity in positive therapeutic outcome to HSCT among individual patients. Our simulations confirmed the clinical observations that termination of cART in the absence of HSCT inevitably leads to viral rebound in all patients ^2^. In addition, we found that extending the duration of cART prior to cessation of therapy only slightly increases the median time to viral rebound and its variance across the four virtual populations, but does not prevent viral rebound across all virtual patients. This suggests that viral rebound after the cessation of cART is an inevitability in the absence of additional interventions, regardless of the duration of antiretroviral treatment. The time to viral rebound after the cessation of cART does differ from patient to patient. This is likely a result of interindividual variability in the underlying infection dynamics and immune response in the patient populations. However, our results indicate that no one mechanistic model parameter singularly determines the time to rebound or the levels of viral load or of T cells. Instead, interactions between multiple model parameters play a complex role in driving treatment responses. Fundamentally, the system itself (and thus the mechanistic model as its surrogate) serves as an integrator that encapsulates the interactions between model parameters, thereby generating the observable outcomes.

Given that cART by itself, regardless of duration, cannot serve as a cure, we added simulation of bone marrow transplant as a potential therapy; specifically, hematopoietic stem cell transplant (HSCT) using cells from a CCR5del32 donor, a mutation which renders those cells resistant to HIV infection. As shown previously using computational modeling, this treatment can result in long-term suppression of HIV that resembles remission or cure, even once cART is stopped. However, the treatment is not predicted to work for all patients. Thus, we explored which mechanistic model parameters would be predictive of therapeutic success, i.e. which parameter values correlated with the therapeutic responders (who are predicted to maintain viral suppression) and non-responders (who are predicted to experience viral rebound). Once again, no one mechanistic parameter or subset of parameter exhibited primary control over the response to treatment. Instead, the mechanistic parameters, with different values from patient to patient, seem to function as a networked whole to collectively determine the response.

In comparison to the mechanistic model parameters, we used PLS-DA to show that virtual clinical observations – for example, viral loads, and T cell levels – are better predictors of therapeutic outcome. These virtual clinical observations were better able to differentiate patients into rebounding and non-rebounding clusters. Moreover, virtual clinical observations are quantities that are more accessible and easily available to be measured from each patient, making them better suited as covariates in combining HSCT with cART as a personalized care for HIV patients. It is likely that these virtual clinical observations are effectively integrators of the underlying mechanistic model, resolving the influence of multiple parameters and providing a readout that is more predictive of therapeutic outcome.

However, there are limitations to our study. While our virtual populations represent the characteristics of the real patient populations well, there is not a direct one-to-one correspondence between our virtual patients and real HIV patients ^8^. Furthermore, the complex virtual clinical trials that we have proposed and simulated are challenging to replicate and validate clinically. The clinical procedures are not trivial, and have generally only been attempted in patients for whom HSCT was already indicated due to concomitant cancer diagnoses. So far, only five patients have been shown to benefit from this HSCT treatment, out of eleven total in whom the procedure has been attempted. Thus, to extrapolate our findings to actual clinical scenarios, one could create patient-specific digital twin virtual populations and perform simulations to confirm our findings. Obtaining sufficient matched donors with CCR5del32 to treat current HIV-infected patients would be extremely difficult, though advances in gene editing technology could enable modification of autologous cells. Because HSCT is an invasive procedure that is primarily reserved for treating hematologic malignancies, considerable evidence would be needed to support replacing long-term cART treatment which such an aggressive treatment. Given the difficulty in validating these simulated observations in human populations, testing of HSCT treatments in nonhuman primates may offer a path to understand the drivers of therapeutic success ^15,16^.

## Acknowledgements

This work was supported by the Rockfish high-performance computing cluster at the Maryland Advanced Research Computing Center (MARCC).

## Author Contributions

J.C. and F.M.G. wrote and revised the manuscript. J.C. and F.M.G. conceptualized the study and designed the experiments. J.C. performed the experiments. J.C. and F.M.G. analyzed and interpreted the experimental results. All authors reviewed and approved the final manuscript.

## Availability of data and materials

The datasets and models for replicating the findings and conclusions of this article are included in the supplementary information section of this article.

## Conflict of Interest

The authors declare no conflict of interest.

## Supplementary Materials for

**Figure S1.**
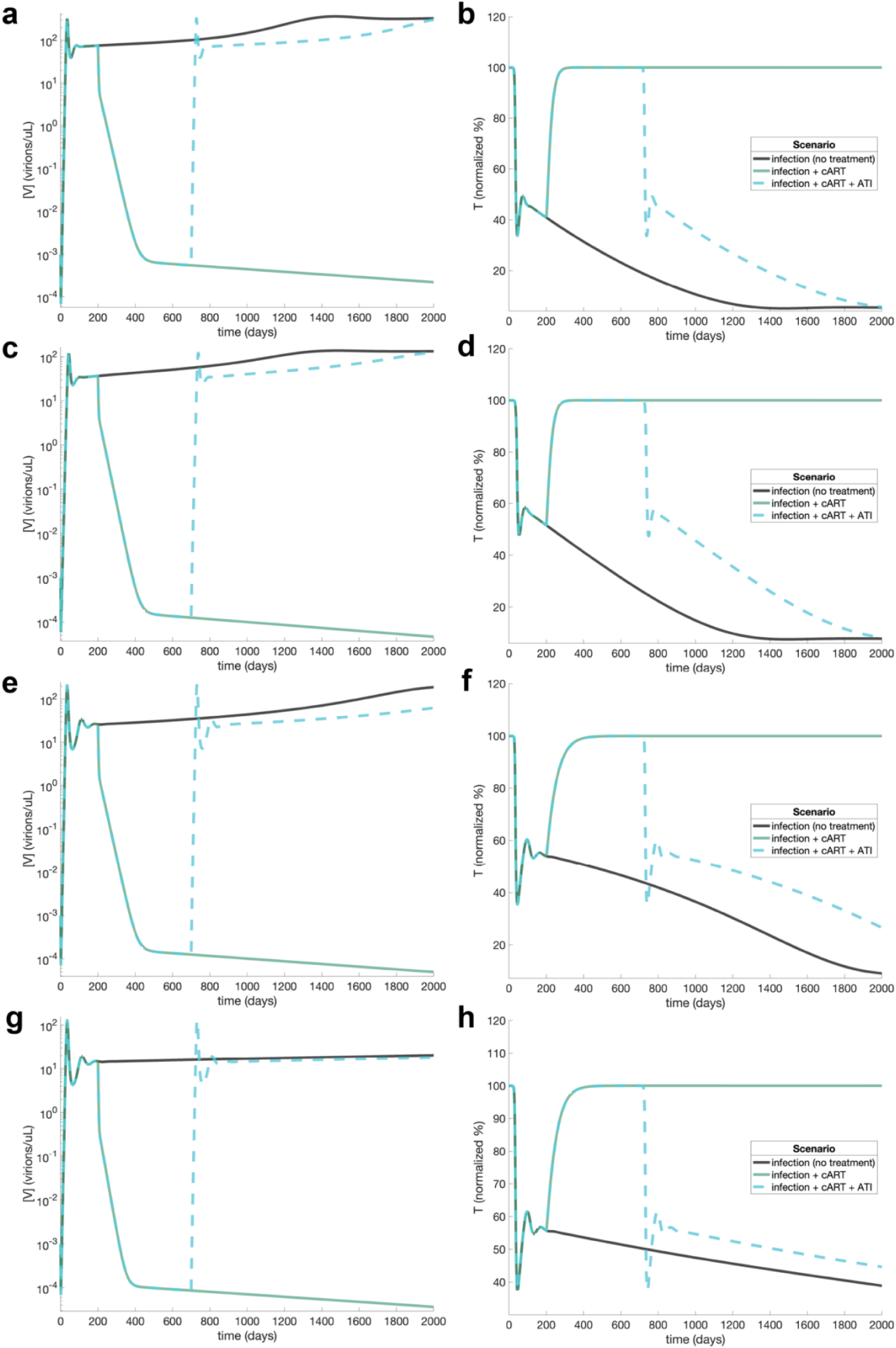
Simulations of infection, combination antiretroviral therapy (cART), and treatment interruption (TI) from mean parameter values and initial conditions for each virtual population. (**a,b**) Population 1; (**c,d**) Population 2; (**e,f**) Population 3; (**g,h**) Population 4. (**a,c,e,g**) Predicted viral loads. (**b,d,f,h**) Predicted CD4+ T cell counts. CD4+ T cell counts were normalized by the number of initial CD4+ T cells present in the system. Three different scenarios are simulated: infection only; infection followed by cART; and infection followed by cART and then TI. The simulated scenarios consist of: infection at day 0; cART from day 200; and TI at day 700.

**Figure S2.**
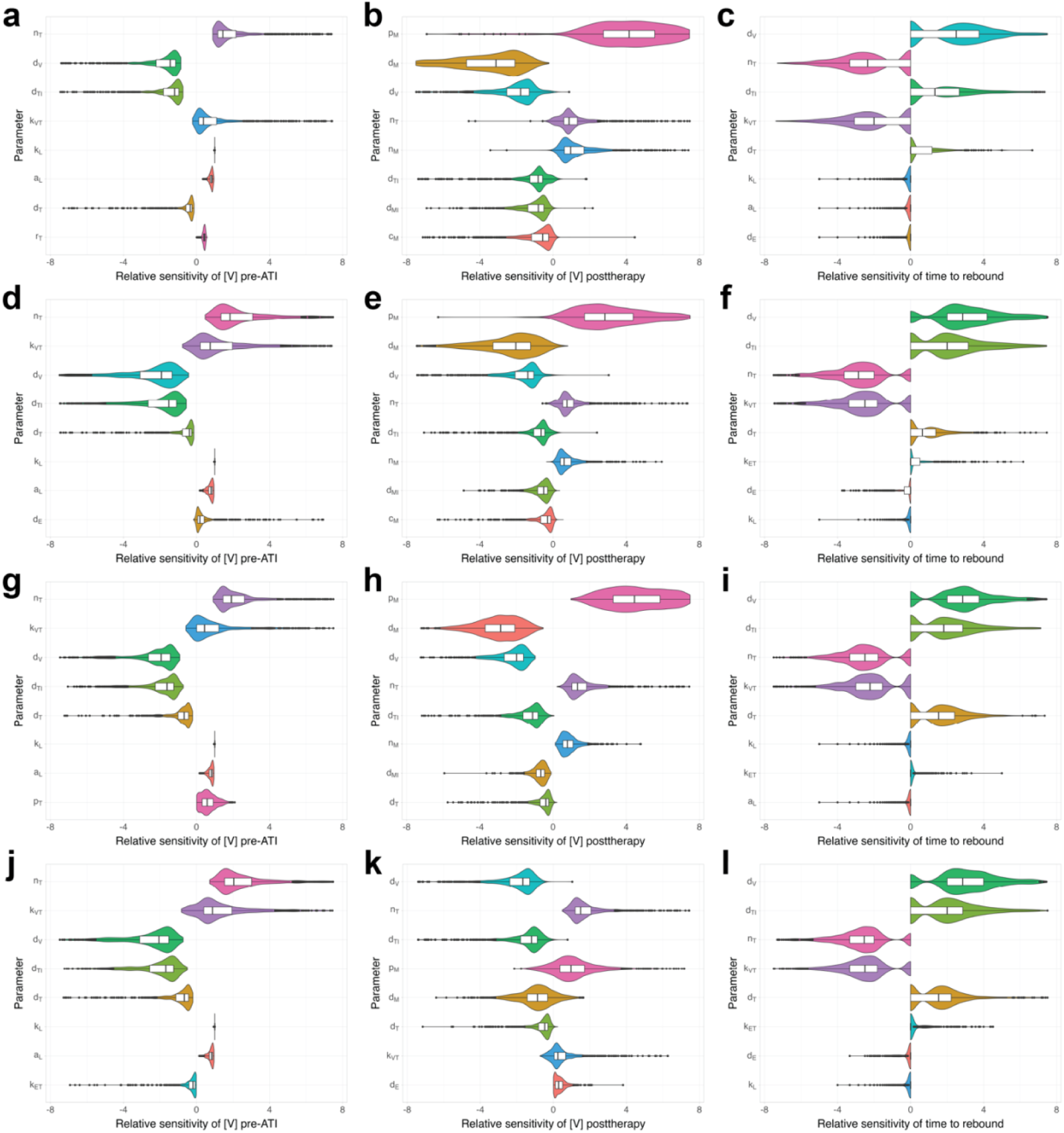
Distribution of local sensitivity of key metrics on model parameters in each virtual population. (**a-c**) Population 1; (**d-f**) Population 2; (**g-i**) Population 3; (**j-l**) Population 4. Relative sensitivity is defined in Figure 5. The violin distributions represent the probability density estimates of relative sensitivity across the virtual patients. Outliers are shown in black dots; the box represents the 25^th^ percentile, median, and 75^th^ percentile. Only parameters with top one-third sensitivity values are shown. Three key metrics are evaluated for each virtual patient: viral load immediately prior to ATI ([V]_pre-ATI_), viral load at day X post-TI ([V]_post-ATI_), and time to viral rebound (T_VR_).

**Figure S3.**
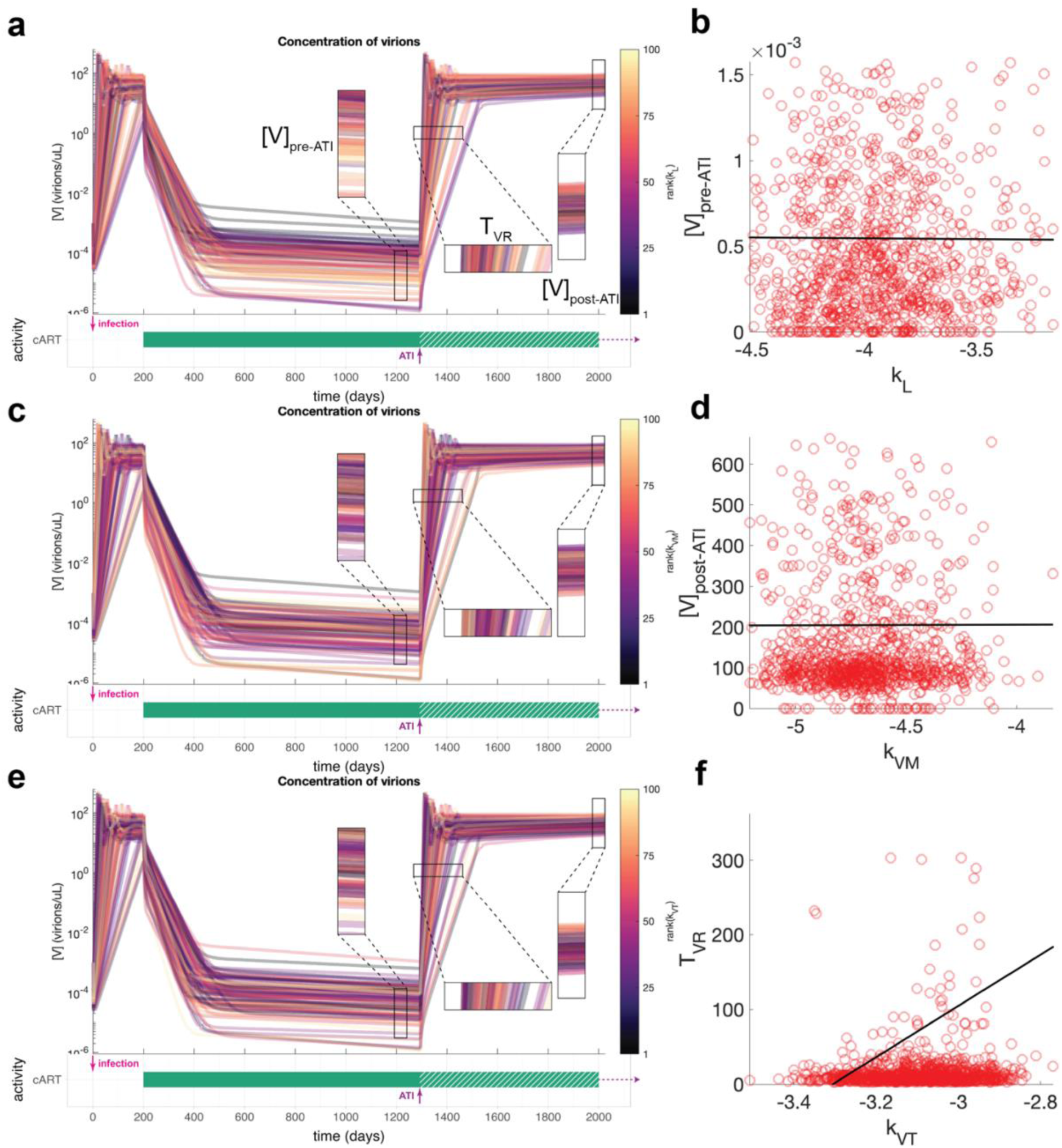
Simulations of infection, combination antiretroviral therapy (cART), and treatment interruption (TI) in the virtual population. (**a,c,e**) Predicted viral loads for 100 virtual patients in Population 2. The virtual clinical trial consists of: infection at day 0; cART at day 200; and TI at day 1300. The individual patient trajectories are colored by the rank of different model parameters to emphasize distinct features of the simulations: (**a**) rate of CD4+ T cell latency conversion (*k_L_*) for viral load immediately prior to ATI ([V]_pre-TI_); (**c**) rate of macrophage infection by virus (*k_VM_*) for viral load at day X post-TI ([V]_post-TI_); and (**e**) rate of CD4+ T cell infection by virus for time to viral rebound (*T_VR_*). These were the top driving model parameters identified for each key metric in previous Sobol global sensitivity analysis. (**b,d,f**) Direct comparison of key metrics with suggested driving model parameters. Each red circle = one virtual patient; black shows line of best fit.

**Figure S4.**
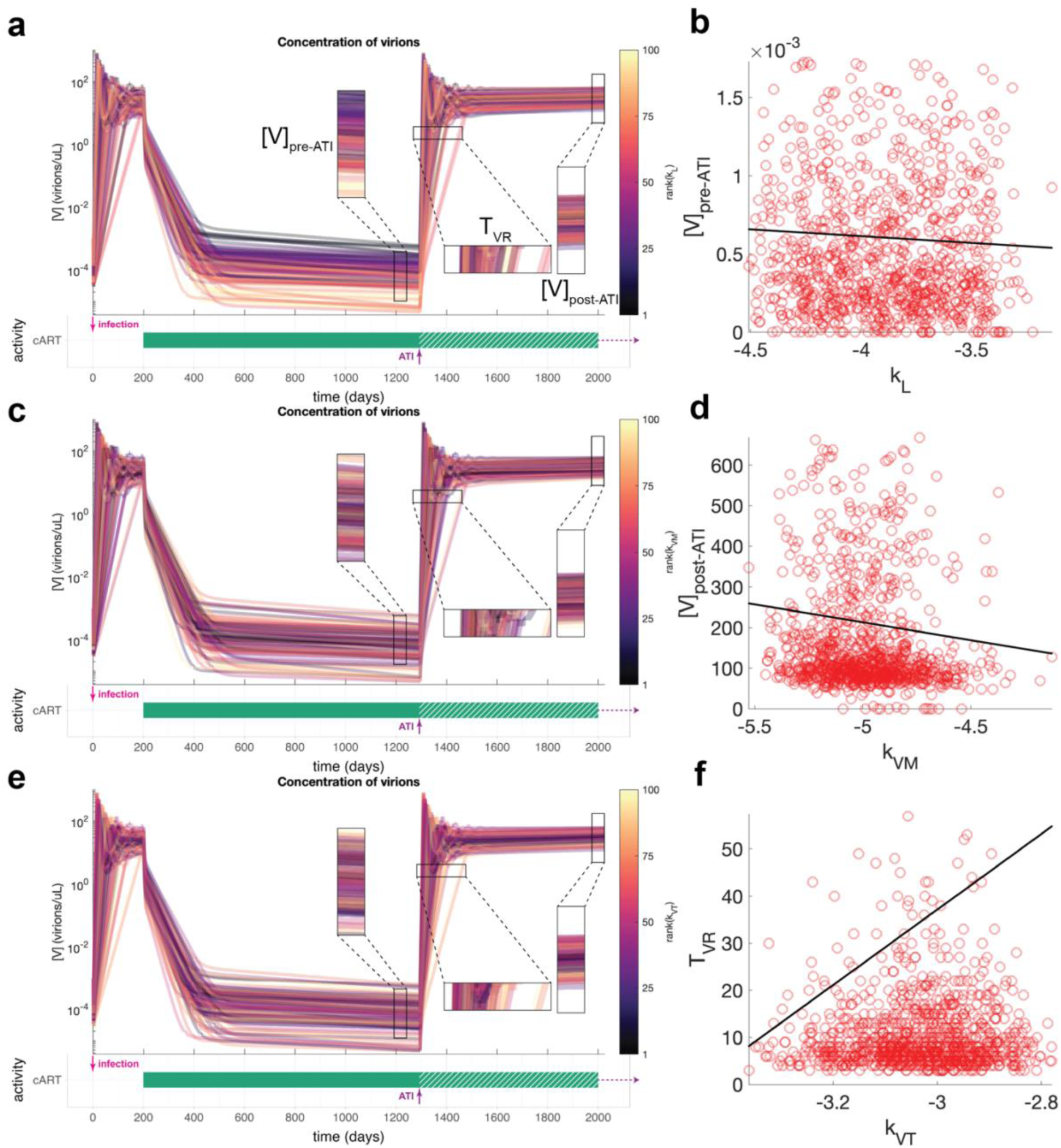
Simulations of infection, combination antiretroviral therapy (cART), and treatment interruption (TI) in the virtual population. (**a,c,e**) Predicted viral loads for 100 virtual patients in Population 3. The virtual clinical trial consists of: infection at day 0; cART at day 200; and TI at day 1300. The individual patient trajectories are colored by the rank of different model parameters to emphasize distinct features of the simulations: (**a**) rate of CD4+ T cell latency conversion (*k_L_*) for viral load immediately prior to TI ([V]_pre-TI_); (**c**) rate of macrophage infection by virus (*k_VM_*) for viral load at day X post-TI ([V]_post-TI_); and (**e**) rate of CD4+ T cell infection by virus for time to viral rebound (*T_VR_*). These were the top driving model parameters identified for each key metric in previous Sobol global sensitivity analysis. (**b,d,f**) Direct comparison of key metrics with suggested driving model parameters. Each red circle = one virtual patient; black shows line of best fit.

**Figure S5.**
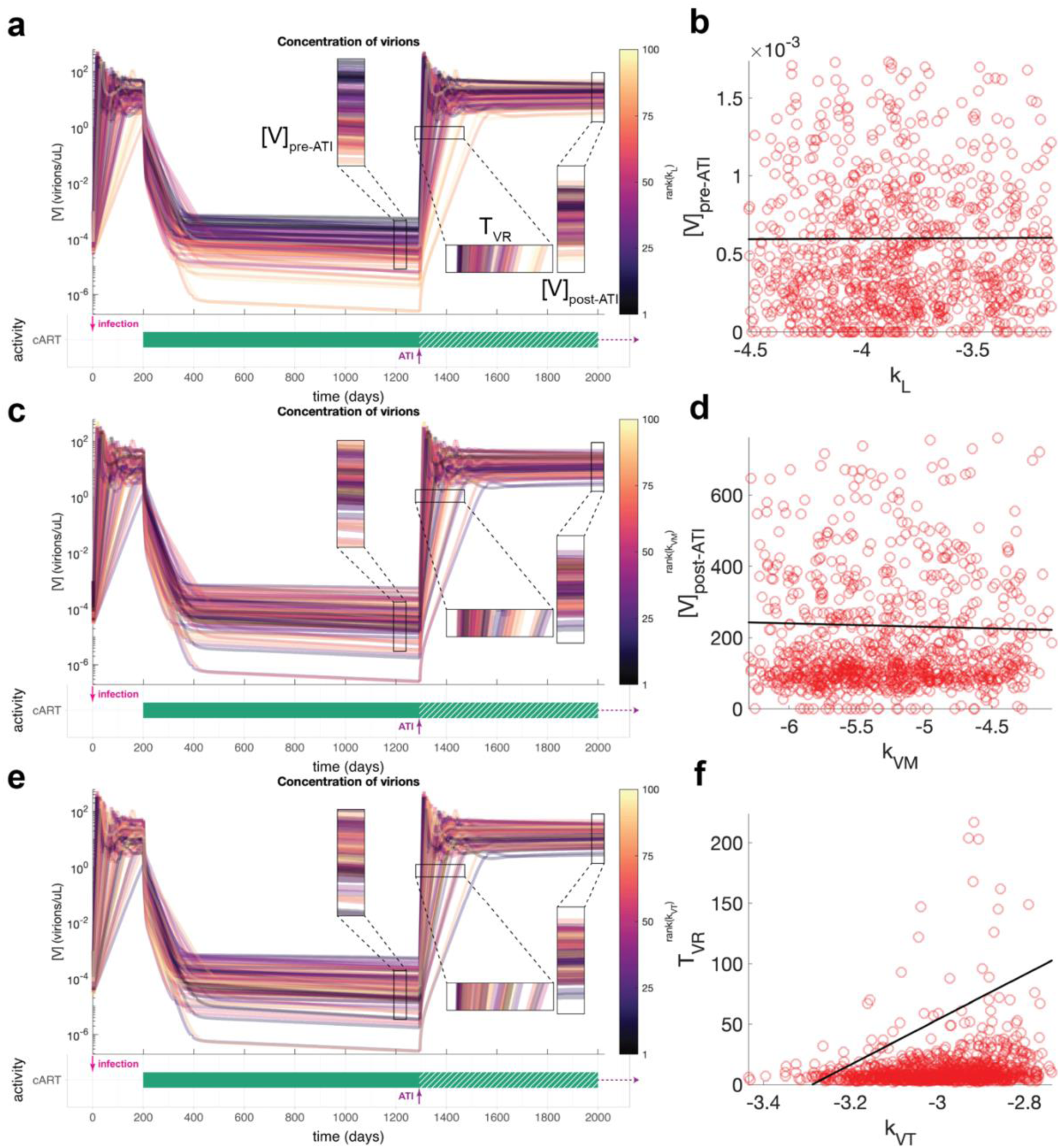
Simulations of infection, combination antiretroviral therapy (cART), and treatment interruption (TI) in the virtual population. (**a,c,e**) Predicted viral loads for 100 virtual patients in Population 4. The virtual clinical trial consists of: infection at day 0; cART at day 200; and TI at day 1300. The individual patient trajectories are colored by the rank of different model parameters to emphasize distinct features of the simulations: (**a**) rate of CD4+ T cell latency conversion (*k_L_*) for viral load immediately prior to TI ([V]_pre-TI_); (**c**) rate of macrophage infection by virus (*k_VM_*) for viral load at day X post-TI ([V]_post-TI_); and (**e**) rate of CD4+ T cell infection by virus for time to viral rebound (*T_VR_*). These were the top driving model parameters identified for each key metric in previous Sobol global sensitivity analysis. (**b,d,f**) Direct comparison of key metrics with suggested driving model parameters. Each red circle = one virtual patient; black shows line of best fit.

**Figure S6.**
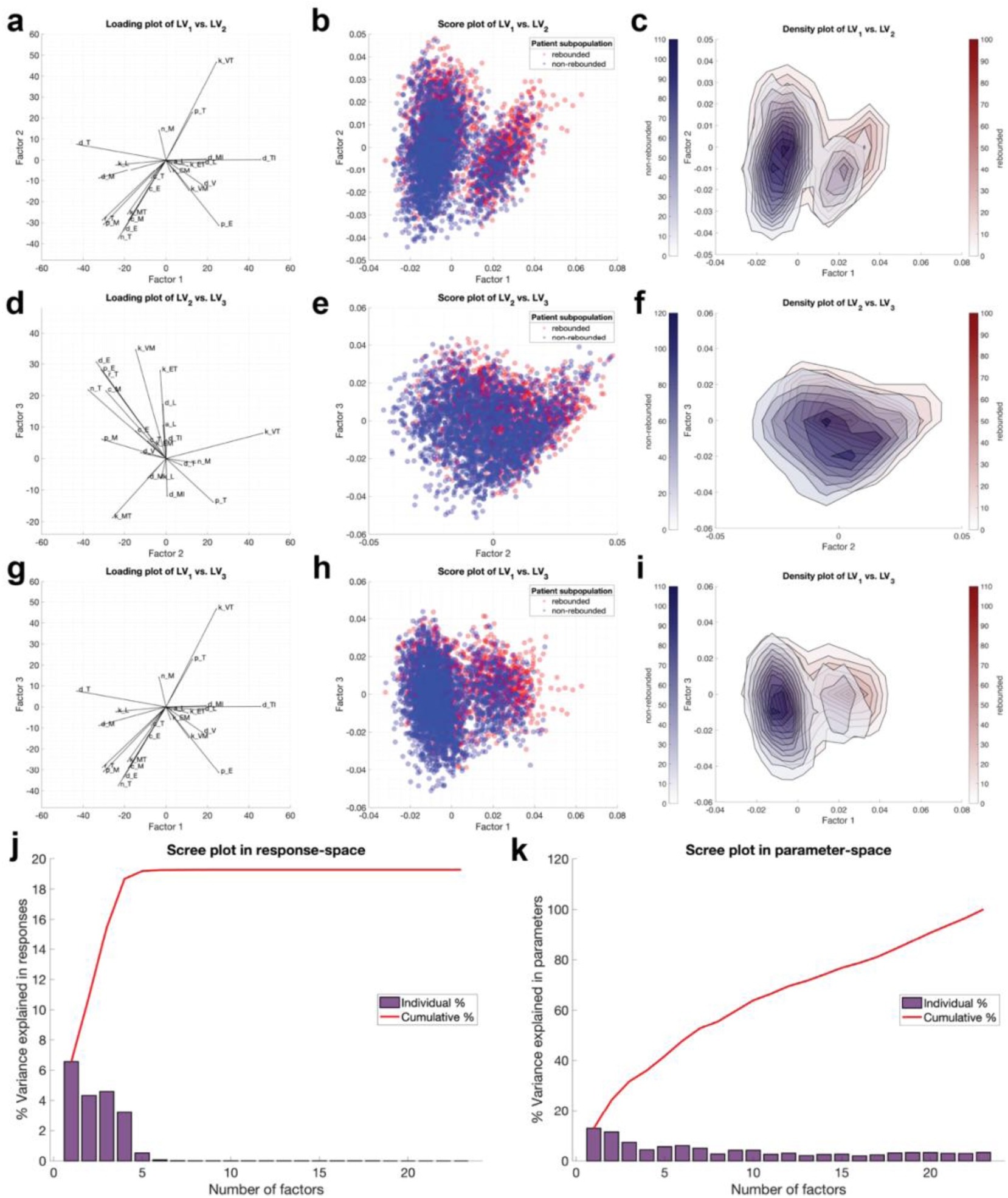
Model parameters as drivers of HSCT therapeutic success of HIV-infected patients in the combined population. This is an extended version of Figure 12. The virtual patients undergoing hematopoietic stem cell transplantation (HSCT) were segregated into responders (no viral rebound, blue) and nonreseponders (viral rebound, red) based on the final viral loads. The threshold for viral rebound is defined by the limit of detection at <50 copies/mL. Partial least squares (PLS) regression used to identify latent variables that best separate the groups. (**a,d,g**) Loading plot of model parameters with respect to two of the first three latent variables. (**b,e,h**) Score plot (one dot = one virtual patient) with respect to two of the first three latent variables. (**c,f,i**) Density plots corresponding to the patient scatter plots shown in the middle column. (**j-k**) Scree plots quantify the percent variance explained by the latent variables individually and cumulatively in the input space (model parameter, **k**) and the response space (viral rebound, **j**).

**Figure S7.**
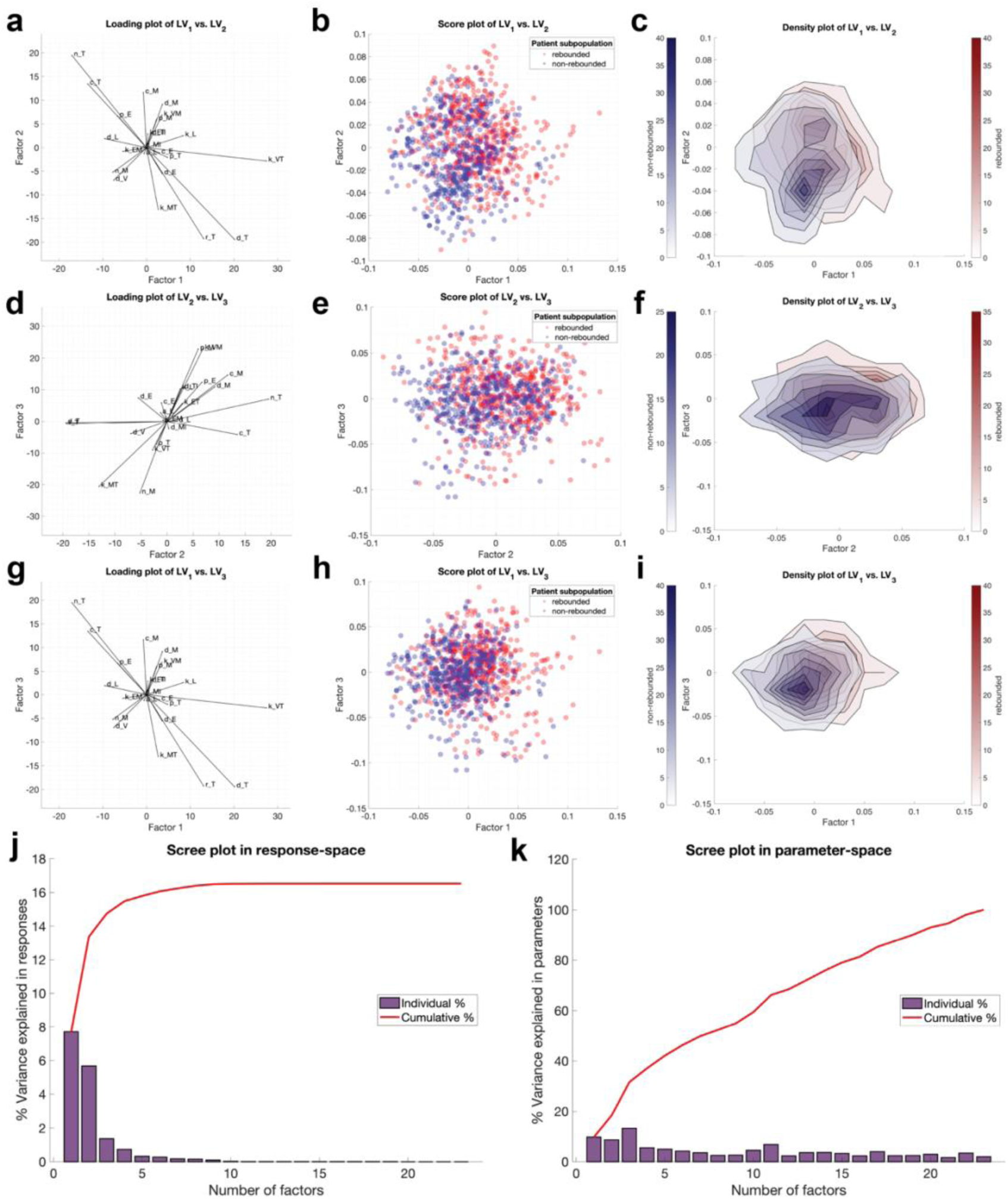
Model parameters as drivers of HSCT therapeutic success of HIV-infected patients in population 1. The virtual patients undergoing hematopoietic stem cell transplantation (HSCT) were segregated into responders (no viral rebound, blue) and nonreseponders (viral rebound, red) based on the final viral loads. The threshold for viral rebound is defined by the limit of detection at <50 copies/mL. Partial least squares (PLS) regression used to identify latent variables that best separate the groups. (**a,d,g**) Loading plot of model parameters with respect to two of the first three latent variables. (**b,e,h**) Score plot (one dot = one virtual patient) with respect to two of the first three latent variables. (**c,f,i**) Density plots corresponding to the patient scatter plots shown in the middle column. (**j-k**) Scree plots quantify the percent variance explained by the latent variables individually and cumulatively in the input space (model parameter, **k**) and the response space (viral rebound, **j**).

**Figure S8.**
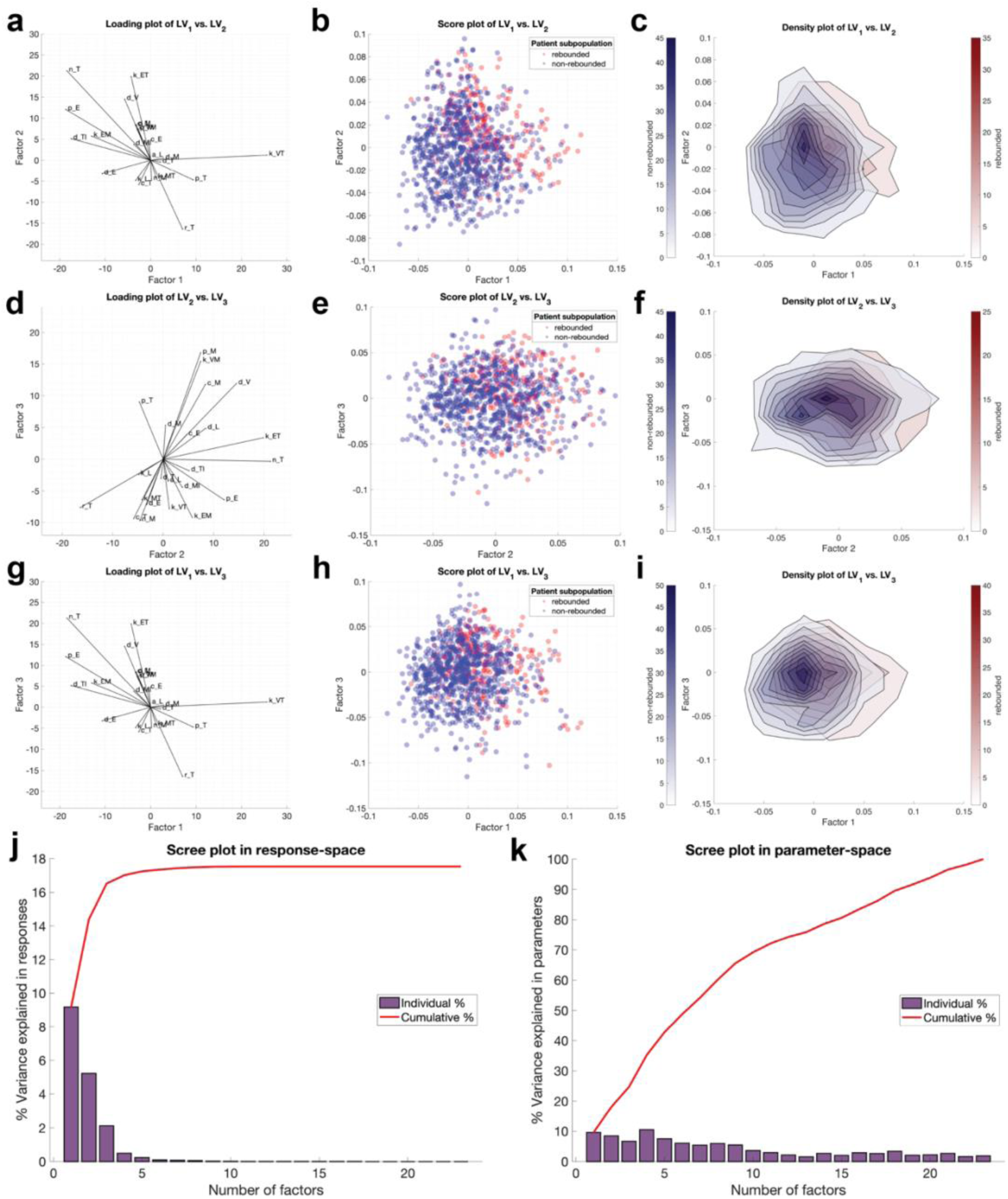
Model parameters as drivers of HSCT therapeutic success of HIV-infected patients in population 2. The virtual patients undergoing hematopoietic stem cell transplantation (HSCT) were segregated into responders (no viral rebound, blue) and nonreseponders (viral rebound, red) based on the final viral loads. The threshold for viral rebound is defined by the limit of detection at <50 copies/mL. Partial least squares (PLS) regression used to identify latent variables that best separate the groups. (**a,d,g**) Loading plot of model parameters with respect to two of the first three latent variables. (**b,e,h**) Score plot (one dot = one virtual patient) with respect to two of the first three latent variables. (**c,f,i**) Density plots corresponding to the patient scatter plots shown in the middle column. (**j-k**) Scree plots quantify the percent variance explained by the latent variables individually and cumulatively in the input space (model parameter, **k**) and the response space (viral rebound, **j**).

**Figure S9.**
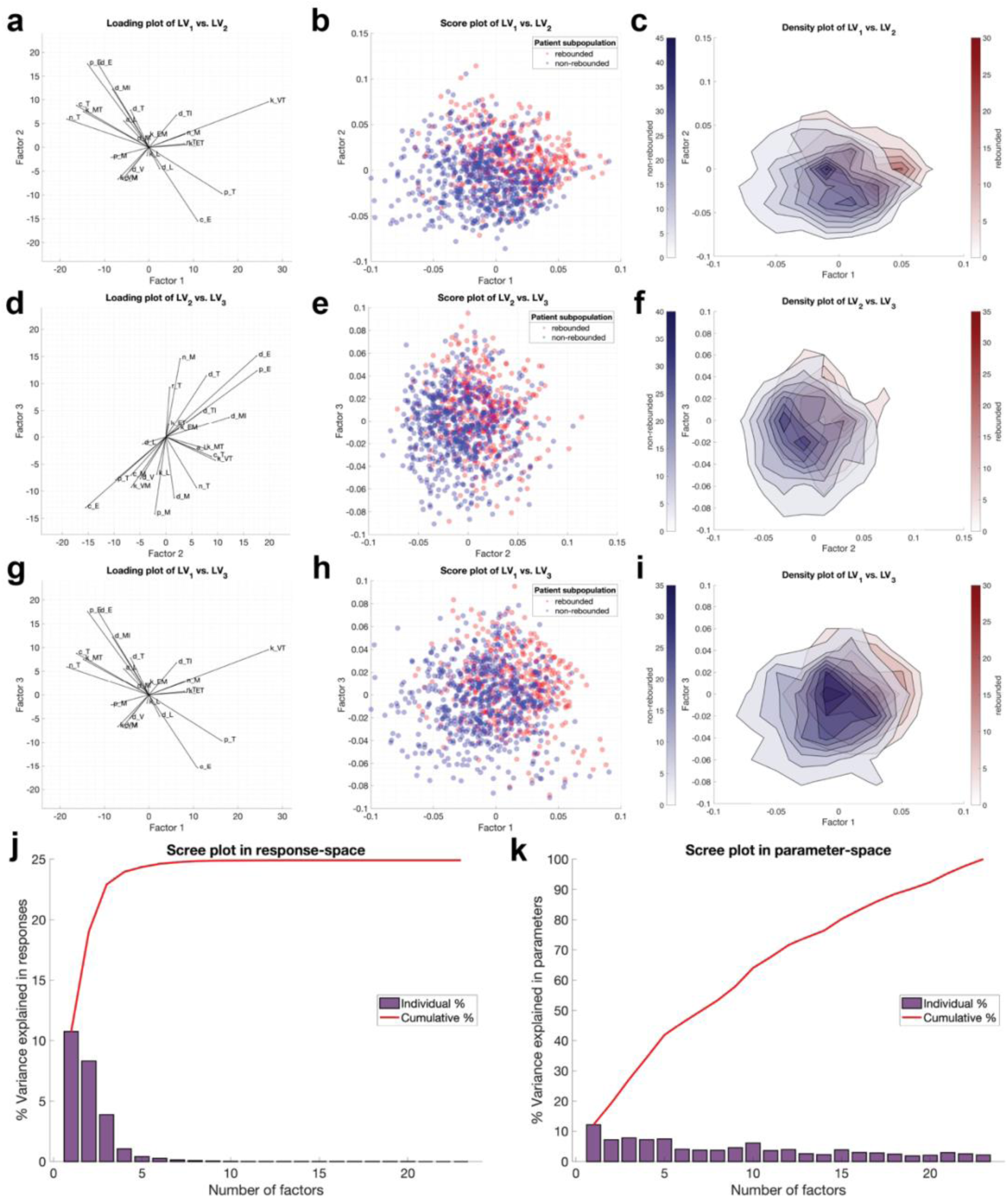
Model parameters as drivers of HSCT therapeutic success of HIV-infected patients in population 3. The virtual patients undergoing hematopoietic stem cell transplantation (HSCT) were segregated into responders (no viral rebound, blue) and nonreseponders (viral rebound, red) based on the final viral loads. The threshold for viral rebound is defined by the limit of detection at <50 copies/mL. Partial least squares (PLS) regression used to identify latent variables that best separate the groups. (**a,d,g**) Loading plot of model parameters with respect to two of the first three latent variables. (**b,e,h**) Score plot (one dot = one virtual patient) with respect to two of the first three latent variables. (**c,f,i**) Density plots corresponding to the patient scatter plots shown in the middle column. (**j-k**) Scree plots quantify the percent variance explained by the latent variables individually and cumulatively in the input space (model parameter, **k**) and the response space (viral rebound, **j**).

**Figure S10.**
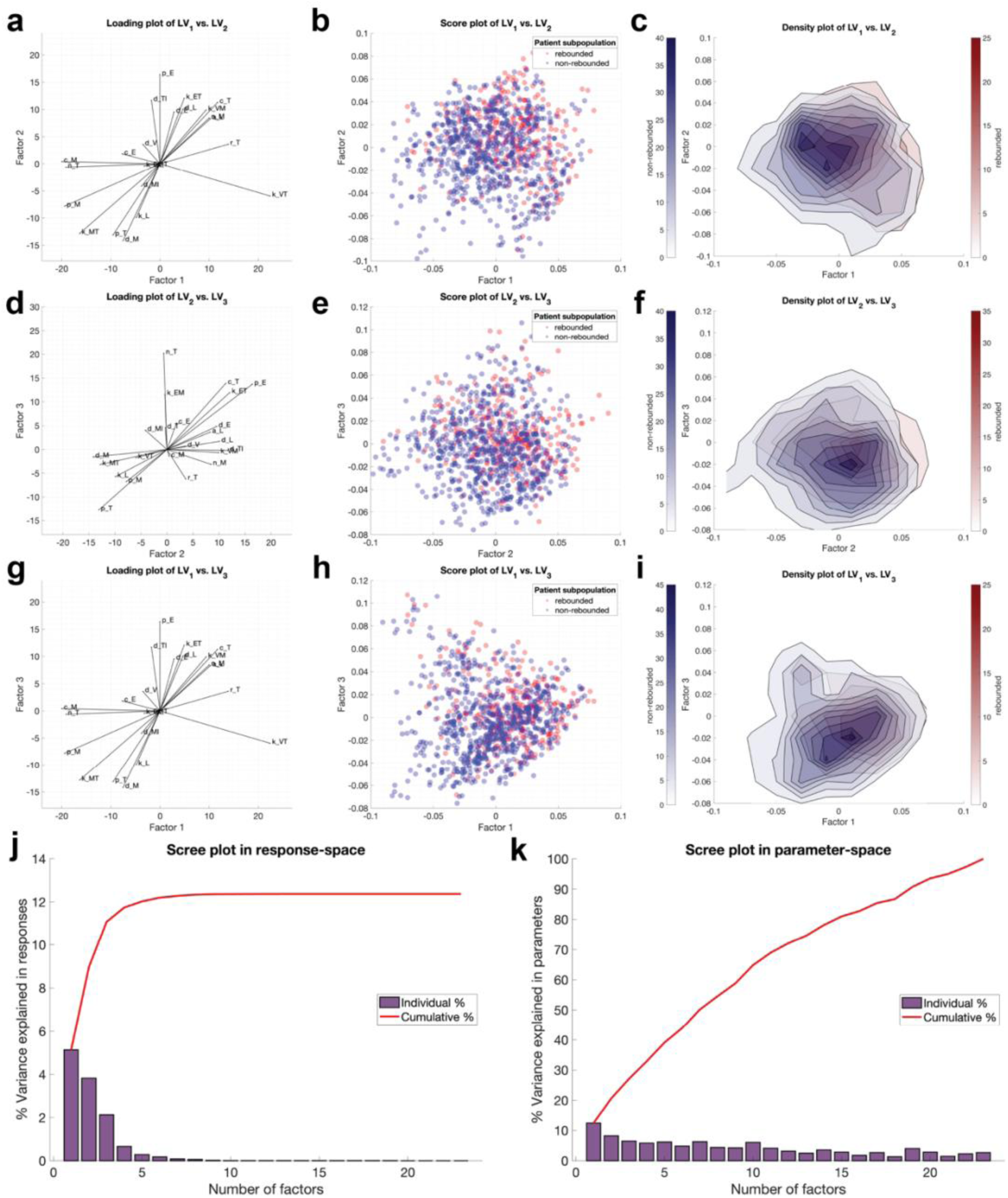
Model parameters as drivers of HSCT therapeutic success of HIV-infected patients in population 4. The virtual patients undergoing hematopoietic stem cell transplantation (HSCT) were segregated into responders (no viral rebound, blue) and nonreseponders (viral rebound, red) based on the final viral loads. The threshold for viral rebound is defined by the limit of detection at <50 copies/mL. Partial least squares (PLS) regression used to identify latent variables that best separate the groups. (**a,d,g**) Loading plot of model parameters with respect to two of the first three latent variables. (**b,e,h**) Score plot (one dot = one virtual patient) with respect to two of the first three latent variables. (**c,f,i**) Density plots corresponding to the patient scatter plots shown in the middle column. (**j-k**) Scree plots quantify the percent variance explained by the latent variables individually and cumulatively in the input space (model parameter, **k**) and the response space (viral rebound, **j**).

**Figure S11.**
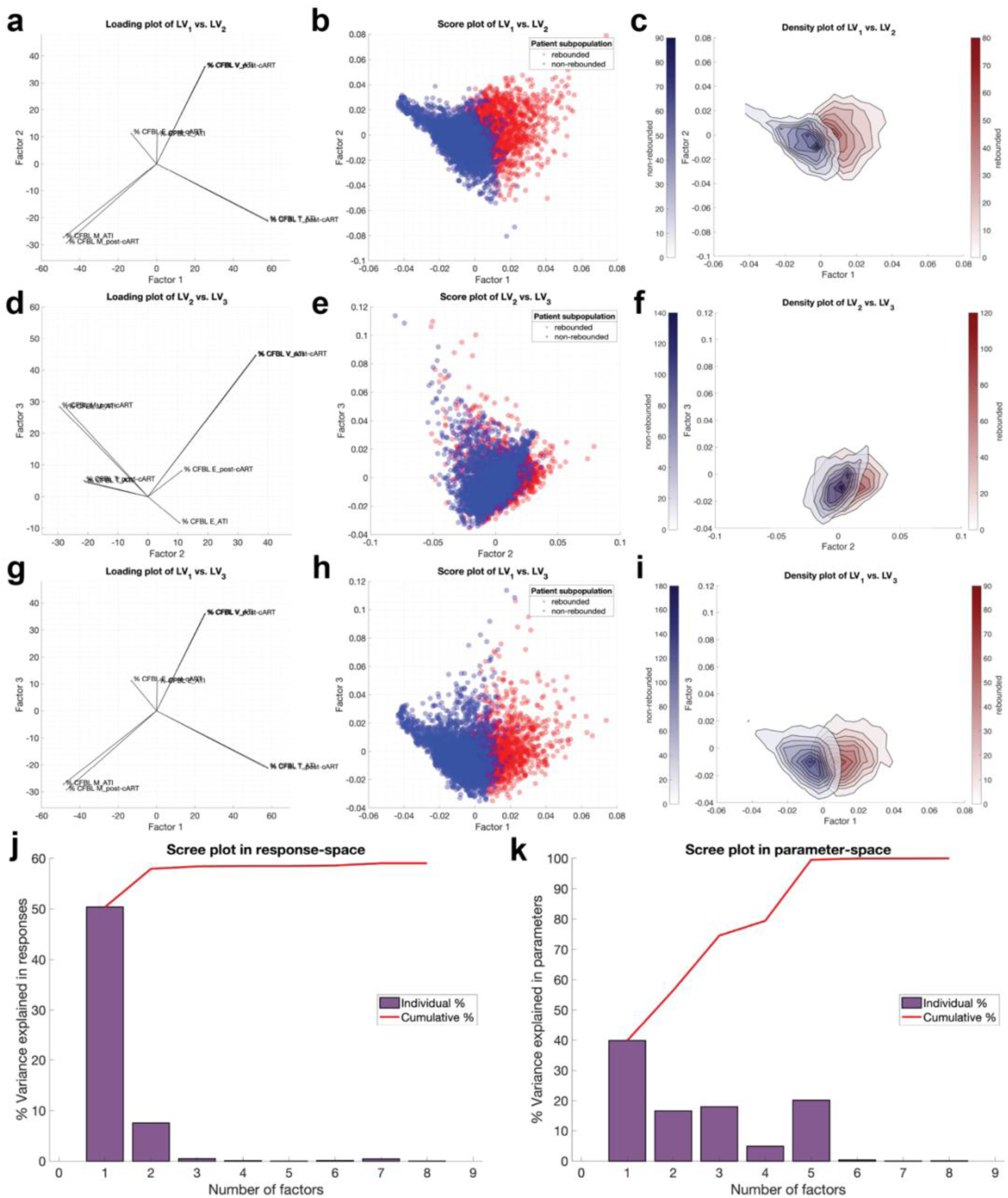
Virtual clinical observations as drivers of HSCT therapeutic success of HIV-infected patients in the combined population. The virtual patients undergoing hematopoietic stem cell transplantation (HSCT) were segregated into responders (no viral rebound, blue) and nonreseponders (viral rebound, red) based on the final viral loads. The threshold for viral rebound is defined by the limit of detection at <50 copies/mL. Partial least squares (PLS) regression used to identify latent variables that best separate the groups. (**a,d,g**) Loading plot of model parameters with respect to two of the first three latent variables. (**b,e,h**) Score plot (one dot = one virtual patient) with respect to two of the first three latent variables. (**c,f,i**) Density plots corresponding to the patient scatter plots shown in the middle column. (**j-k**) Scree plots quantify the percent variance explained by the latent variables individually and cumulatively in the input space (virtual clinical observation, **k**) and the response space (viral rebound, **j**).

**Figure S12.**
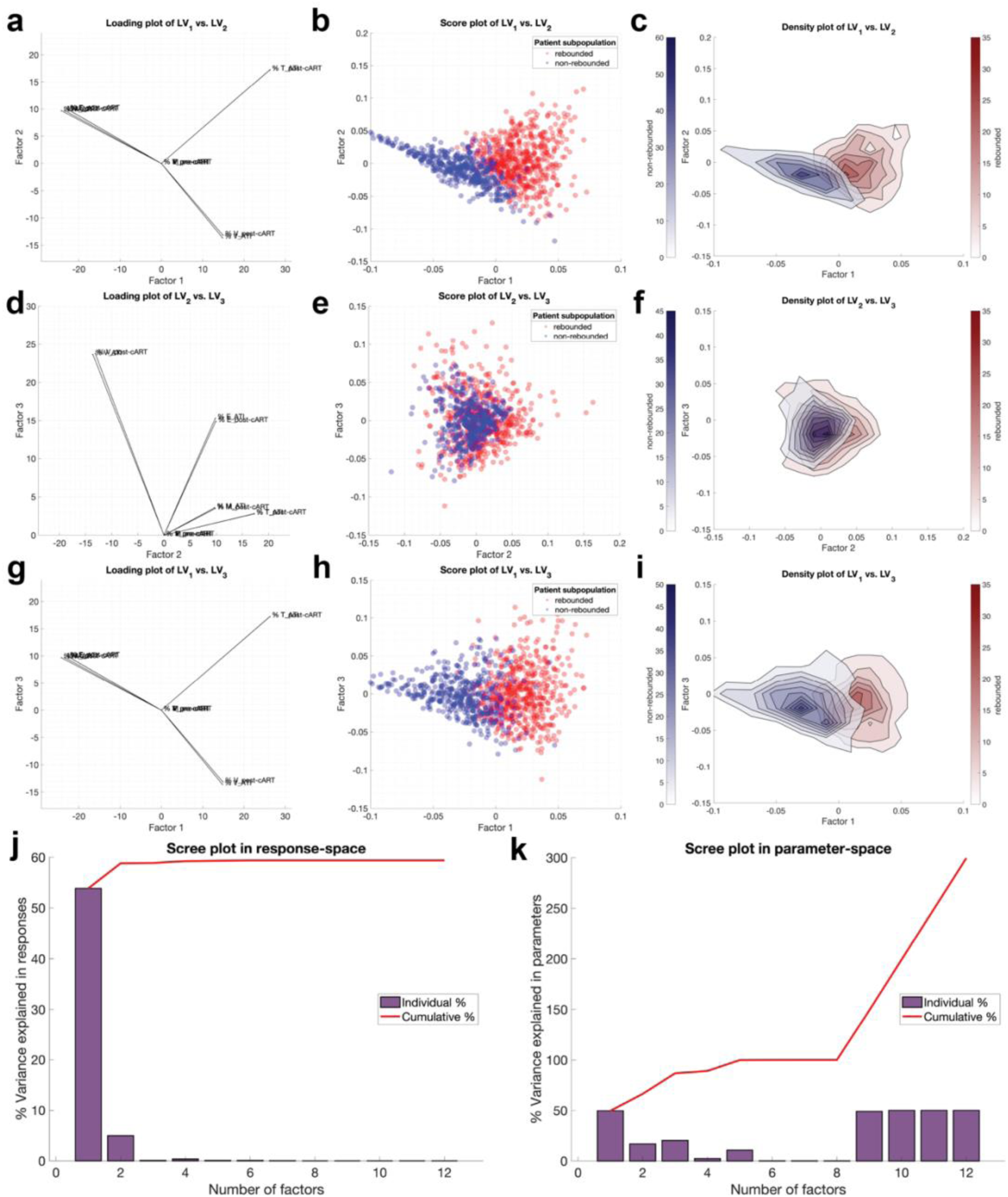
Virtual clinical observations as drivers of HSCT therapeutic success of HIV-infected patients in population 1. The virtual patients undergoing hematopoietic stem cell transplantation (HSCT) were segregated into responders (no viral rebound, blue) and nonreseponders (viral rebound, red) based on the final viral loads. The threshold for viral rebound is defined by the limit of detection at <50 copies/mL. Partial least squares (PLS) regression used to identify latent variables that best separate the groups. (**a,d,g**) Loading plot of model parameters with respect to two of the first three latent variables. (**b,e,h**) Score plot (one dot = one virtual patient) with respect to two of the first three latent variables. (**c,f,i**) Density plots corresponding to the patient scatter plots shown in the middle column. (**j-k**) Scree plots quantify the percent variance explained by the latent variables individually and cumulatively in the input space (virtual clinical observation, **k**) and the response space (viral rebound, **j**).

**Figure S13.**
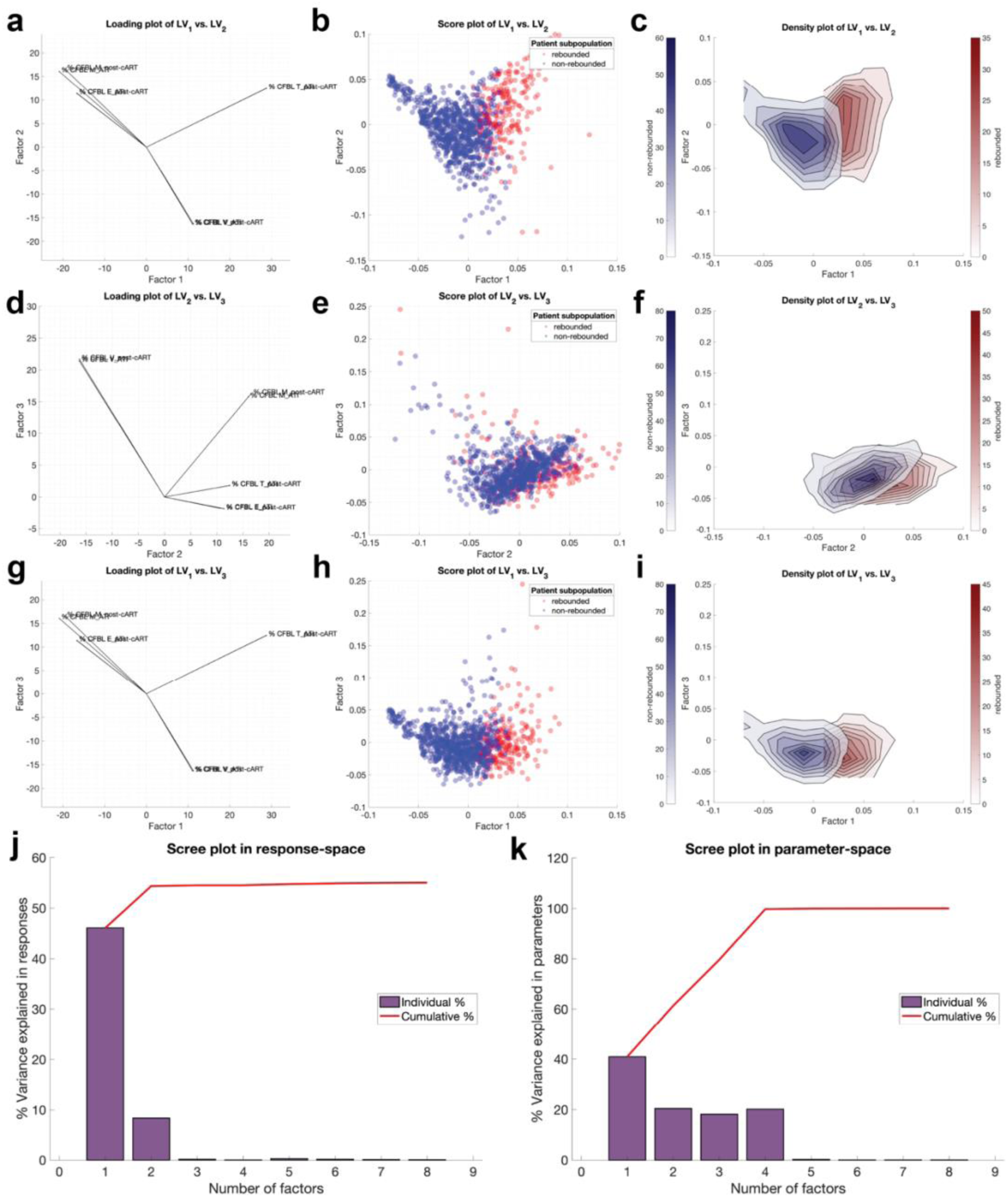
Virtual clinical observations as drivers of HSCT therapeutic success of HIV-infected patients in population 2. The virtual patients undergoing hematopoietic stem cell transplantation (HSCT) were segregated into responders (no viral rebound, blue) and nonreseponders (viral rebound, red) based on the final viral loads. The threshold for viral rebound is defined by the limit of detection at <50 copies/mL. Partial least squares (PLS) regression used to identify latent variables that best separate the groups. (**a,d,g**) Loading plot of model parameters with respect to two of the first three latent variables. (**b,e,h**) Score plot (one dot = one virtual patient) with respect to two of the first three latent variables. (**c,f,i**) Density plots corresponding to the patient scatter plots shown in the middle column. (**j-k**) Scree plots quantify the percent variance explained by the latent variables individually and cumulatively in the input space (virtual clinical observation, **k**) and the response space (viral rebound, **j**).

**Figure S14.**
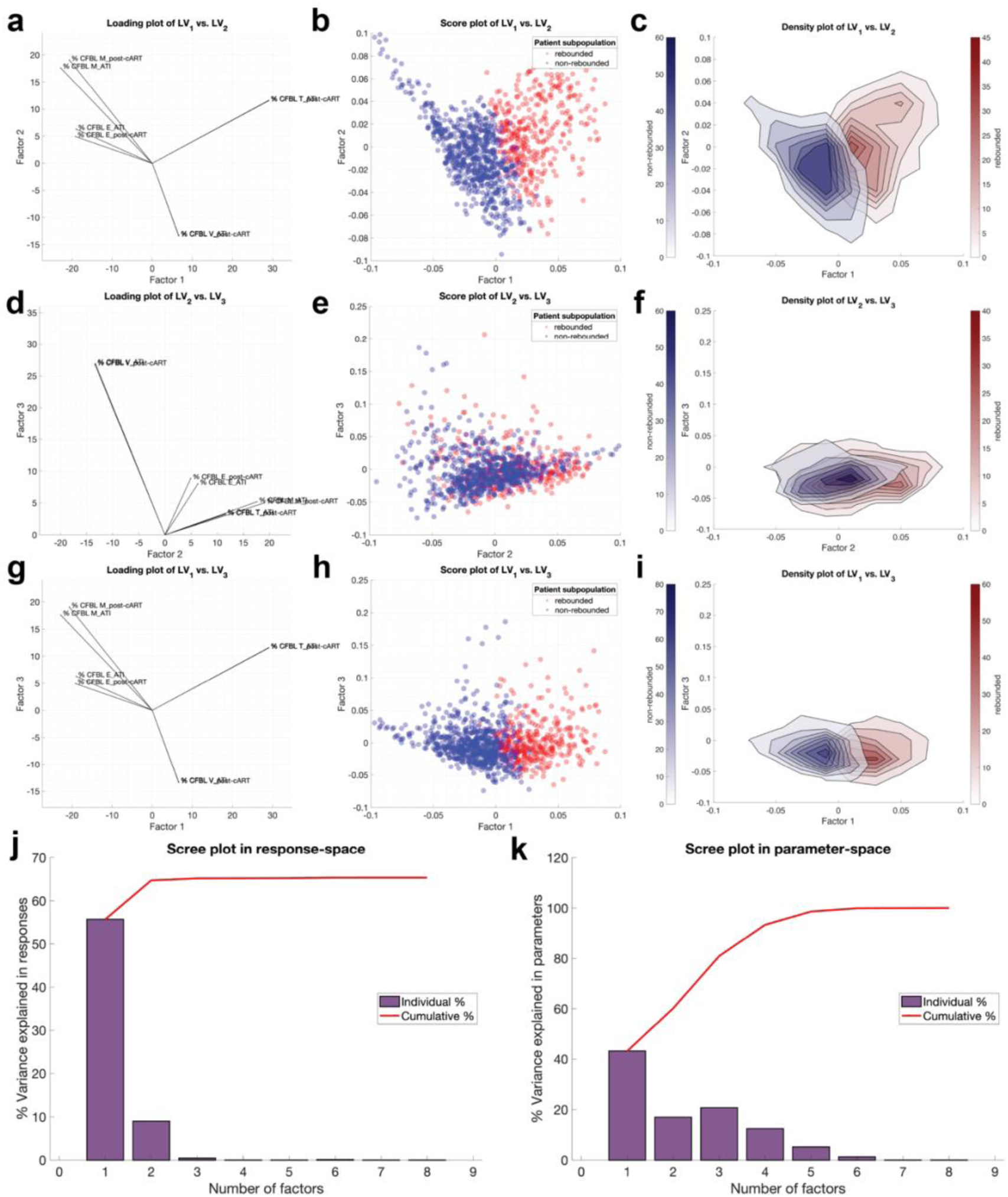
Virtual clinical observations as drivers of HSCT therapeutic success of HIV-infected patients in population 3. The virtual patients undergoing hematopoietic stem cell transplantation (HSCT) were segregated into responders (no viral rebound, blue) and nonreseponders (viral rebound, red) based on the final viral loads. The threshold for viral rebound is defined by the limit of detection at <50 copies/mL. Partial least squares (PLS) regression used to identify latent variables that best separate the groups. (**a,d,g**) Loading plot of model parameters with respect to two of the first three latent variables. (**b,e,h**) Score plot (one dot = one virtual patient) with respect to two of the first three latent variables. (**c,f,i**) Density plots corresponding to the patient scatter plots shown in the middle column. (**j-k**) Scree plots quantify the percent variance explained by the latent variables individually and cumulatively in the input space (virtual clinical observation, **k**) and the response space (viral rebound, **j**).

**Figure S15.**
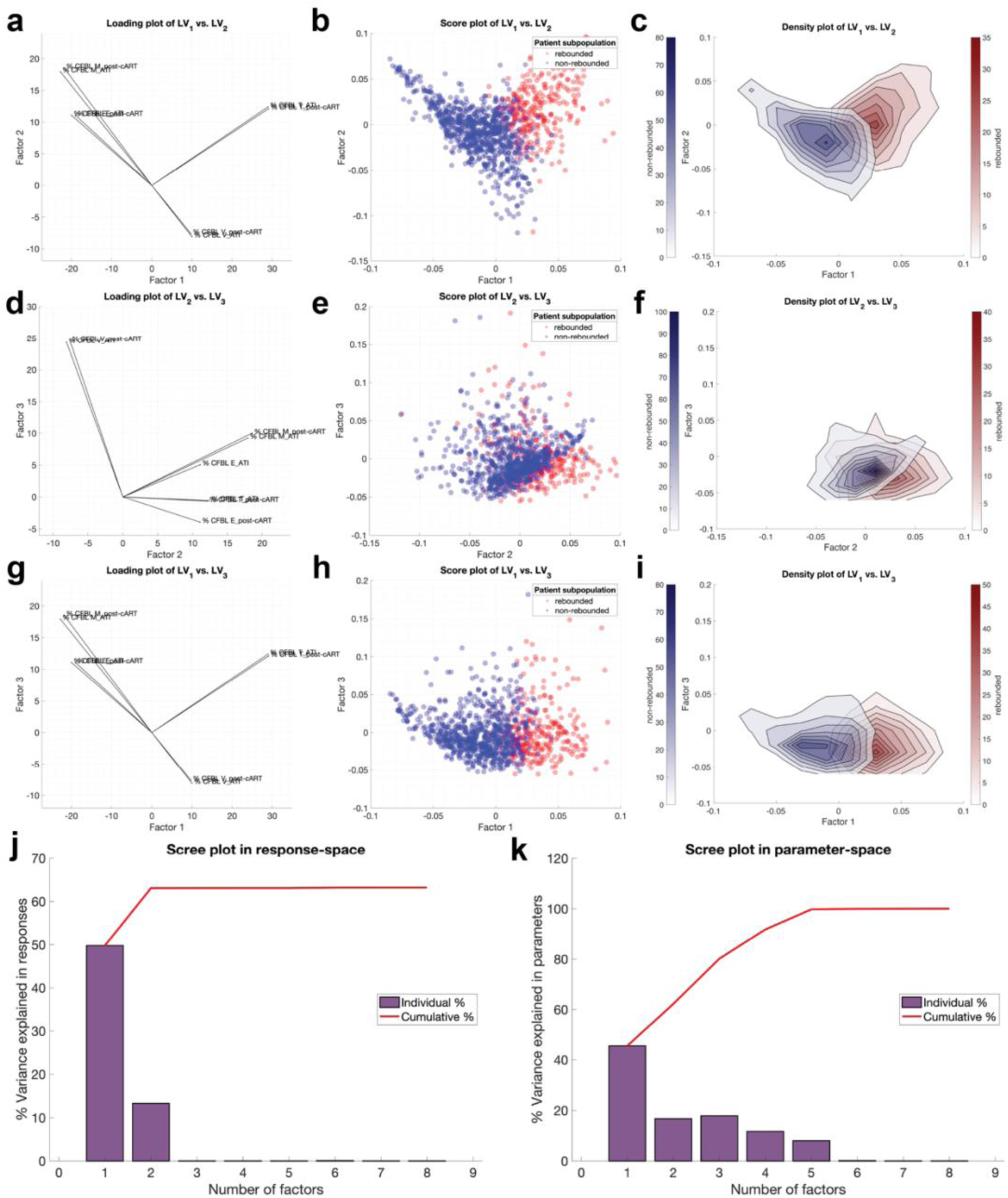
Virtual clinical observations as drivers of HSCT therapeutic success of HIV-infected patients in population 4. The virtual patients undergoing hematopoietic stem cell transplantation (HSCT) were segregated into responders (no viral rebound, blue) and nonreseponders (viral rebound, red) based on the final viral loads. The threshold for viral rebound is defined by the limit of detection at <50 copies/mL. Partial least squares (PLS) regression used to identify latent variables that best separate the groups. (**a,d,g**) Loading plot of model parameters with respect to two of the first three latent variables. (**b,e,h**) Score plot (one dot = one virtual patient) with respect to two of the first three latent variables. (**c,f,i**) Density plots corresponding to the patient scatter plots shown in the middle column. (**j-k**) Scree plots quantify the percent variance explained by the latent variables individually and cumulatively in the input space (virtual clinical observation, **k**) and the response space (viral rebound, **j**).

## Notes

### Competing Interest Statement

The authors have declared no competing interest.

